# Mutant p53 triggers a dynamin-1/APPL1 endosome feedback loop that regulates β1 integrin recycling and migration

**DOI:** 10.1101/408815

**Authors:** Ashley M. Lakoduk, Philippe Roudot, Marcel Mettlen, Heather M. Grossman, Sandra L. Schmid, Ping-Hung Chen

## Abstract

Multiple mechanisms contribute to cancer cell progression and metastatic activity, including changes in endocytic trafficking and signaling of cell surface receptors. We report that gain-of-function (GOF) mutant p53 expression enhances β integrin and EGF receptor recycling and increases cell migration by triggering a positive feedback loop involving the activation of dynamin-1 (Dyn1) and accumulation of a spatially-restricted subpopulation of APPL1-positive ‘perimeter’ endosomes. DNM1 is upregulated at both the mRNA and protein levels in a manner dependent on expression of GOF mutant p53. Perimeter APPL1 endosomes are required for rapid recycling of EGFR and β1 integrins and modulate Akt signaling and Dyn1 activation to create the positive feedback loop that culminates in increased focal adhesion turnover and cell migration. Thus, Dyn1- and Akt-dependent perimeter APPL1 endosomes function as a nexus, integrating signaling and receptor trafficking, that can be co-opted by cancer cells for mutant p53-driven migration and invasion.

## Introduction

Cancer cell invasion and metastasis are mechanistically heterogeneous and adaptive processes contributing to the majority of cancer-related deaths (Bacac and Stamenkovic, 2008; Friedl and Alexander, 2011). While driver gene mutations and epigenetic alterations can increase cancer cell proliferation, survival, invasion and migration, they cannot account for all of the metastatic traits acquired through evolution of aggressive cancer cells (Schmid, 2017). The underlying mechanisms governing the plasticity in transition from primary to aggressive tumors, during which cancer cells acquire their adaptive metastatic abilities, remain largely unknown. Understanding the possible mechanisms leading to cancer metastasis is crucial for successful cancer treatment.

Signaling receptors, including receptor tyrosine kinases (RTKs) and integrins, control many aspects of cell physiology and behavior and are frequently dysregulated in and associated with the initiation and progression of cancer (Sever and Brugge, 2015). Their signaling activities are, in turn, modulated by endocytic trafficking (Mellman and Yarden, 2013). Indeed, gain-of-function (GOF) p53 mutations, which contribute to a more invasive phenotype in multiple cancers (Lang et al., 2004; Olive et al., 2004) can trigger increased recycling of EGFR, cMET and β1 integrins (Lanzetti and Di Fiore, 2017; Muller et al., 2009; Muller et al., 2013). This, in turn, results in increased invasion and migration. The mechanisms responsible for GOF p53 mutant-dependent changes in endocytic trafficking remain poorly understood.

Endocytic trafficking of signaling receptors, which are internalized primarily via clathrin-mediated endocytosis (CME), involves delivery through distinct early endosomal compartments marked by the scaffold proteins APPL1 (adaptor protein, phosphotyrosine interacting with PH domain and leucine zipper 1) and EEA1 (early endosome antigen 1) (Kalaidzidis et al., 2015; Zoncu et al., 2009). Receptors can be recycled back to the cell surface along either fast (i.e. directly from early endosomes) or slow (i.e. via perinuclear recycling endosomes) pathways. Alternatively, receptors can be packaged in intralumenal vesicles and delivered to lysosomes for degradation (Kalaidzidis et al., 2015). Difficulties in quantitatively measuring fast recycling render this the least mechanistically understood of these trafficking pathways.

GOF p53-dependent receptor recycling requires the Rab11 effector, Rab-coupling protein (RCP) (Muller et al., 2009; Muller et al., 2013). However, RCP expression levels are not regulated by p53, thus whether mutant p53 expression directly or indirectly alters expression of components of the endocytic machinery to altered endocytic trafficking remains unknown. Also unknown are the identities of the endosomal compartments from which this recycling occurs, although Rab11 is associated with recycling endosomes and the slow recycling pathway (Wandinger-Ness and Zerial, 2014).

The temporal and functional relationships between the early APPL1 and EEA1 endosomes also remain incompletely defined (Kalaidzidis et al., 2015; Zoncu et al., 2009). APPL1-positive endosomes are often referred to as ‘signaling endosomes’ because APPL1, through its scaffolding properties, regulates many signaling events including Akt/GSK3β activity (Diggins and Webb, 2017; Ding et al., 2016; Schenck et al., 2008). In addition, APPL1 endosomes have been linked to the regulation of cell migration (Broussard et al., 2012; Ding et al., 2016; Tan et al., 2010) and to recycling of some GPCRs (Jean-Alphonse et al., 2014; Sposini et al., 2017). APPL1 endosomes have been reported to be regulated by PKA signaling downstream of GPCRs (Sposini et al., 2017) and by CME itself (Zoncu et al., 2009). Thus, while still poorly defined, APPL1 endosomes are emerging as important integrators of signaling and endocytic trafficking.

The large GTPase dynamin (Dyn) plays an important role in endocytosis. Vertebrates encode three differentially-expressed isoforms: of these, Dyn2 is uniformly expressed, Dyn1 is highly enriched in neurons and Dyn3 is primarily detected in neurons, testes and lung. We recently reported that the neuronally-enriched and typically quiescent Dyn1 is specifically upregulated and/or activated in many non-small cell lung cancer (NSCLC) lines (Reis et al., 2017; Schmid, 2017). Indeed, Dyn1 has emerged as a nexus in regulating signaling and endocytic trafficking in cancer cells (Chen et al., 2017; Reis et al., 2015; Srinivasan et al., 2018). Its activation leads to altered EGFR trafficking, increased Akt signaling and the accumulation of peripheral APPL1-positive endosomes (hereafter called APPL1 endosomes) (Chen et al., 2017). Together these effects are associated with increased metastatic activity of H1299 NSCLC cells in a mouse model for lung metastasis (Chen et al., 2017). Whether this pathway is also linked to GOF p53-dependent changes in endocytic trafficking is unknown.

We have developed quantitative approaches to study rapid recycling from early endosomes and APPL1 endosome regulation. Applying these tools to study mutant p53-driven alterations in endocytic membrane trafficking and cell migration, we find that both DNM1 and Myosin VI (Myo6) are upregulated in cells expressing mutant p53 and are required for enhanced recruitment of APPL1 to a spatially-restricted subpopulation of endosomes that is also dependent on EGFR and Akt signaling. This mutant p53-triggered and EGFR/Akt/Dyn1-dependent subpopulation of APPL1-positive endosomes functions as a hub to modulate signaling and fast recycling of β1 integrins and enhance the migratory and invasive phenotypes in cancer cells. This study reveals mechanisms that govern complex reciprocal regulation between endocytic trafficking and receptor signaling adapted by cancer cells to acquire properties that can enhance their metastatic activities during cancer progression.

## Results

### EGFR/Akt signaling regulates a subpopulation of peripheral APPL1 endosomes in p53-driven cancer cells

We sought to identify the underlying mechanisms and pathways by which the p53 gain-of-function mutant, R273H, drives alterations in the recycling and signaling of surface receptors and cancer invasion (Muller et al., 2009). Given the recently discovered link between EGFR/Akt signaling, Dyn1 activation and the accumulation of APPL1 endosomes (Chen et al., 2017; Reis et al., 2015; Srinivasan et al., 2018), we tested whether these factors might also be required for mutant p53-driven changes in early endocytic trafficking.

H1975 NSCLC cells, which carry a homozygous R273H GOF p53 mutation, were serum starved and then stimulated with 20 ng/ml of EGF. The distribution of APPL1 endosomes was then determined by immunofluorescence. APPL2, its closely-related isoform, was also examined to determine the specificity of these interactions (Figures 1A). To reduce noise from out-of-focus planes and to limit our analyses to peripherally-localized early endosomes, total internal reflection fluorescence (TIRF) microscopy was used after adjusting the angle of illumination to generate a theoretical penetration depth of ∼200nm (i.e. “thick TIRF”).

**Figure 1.**
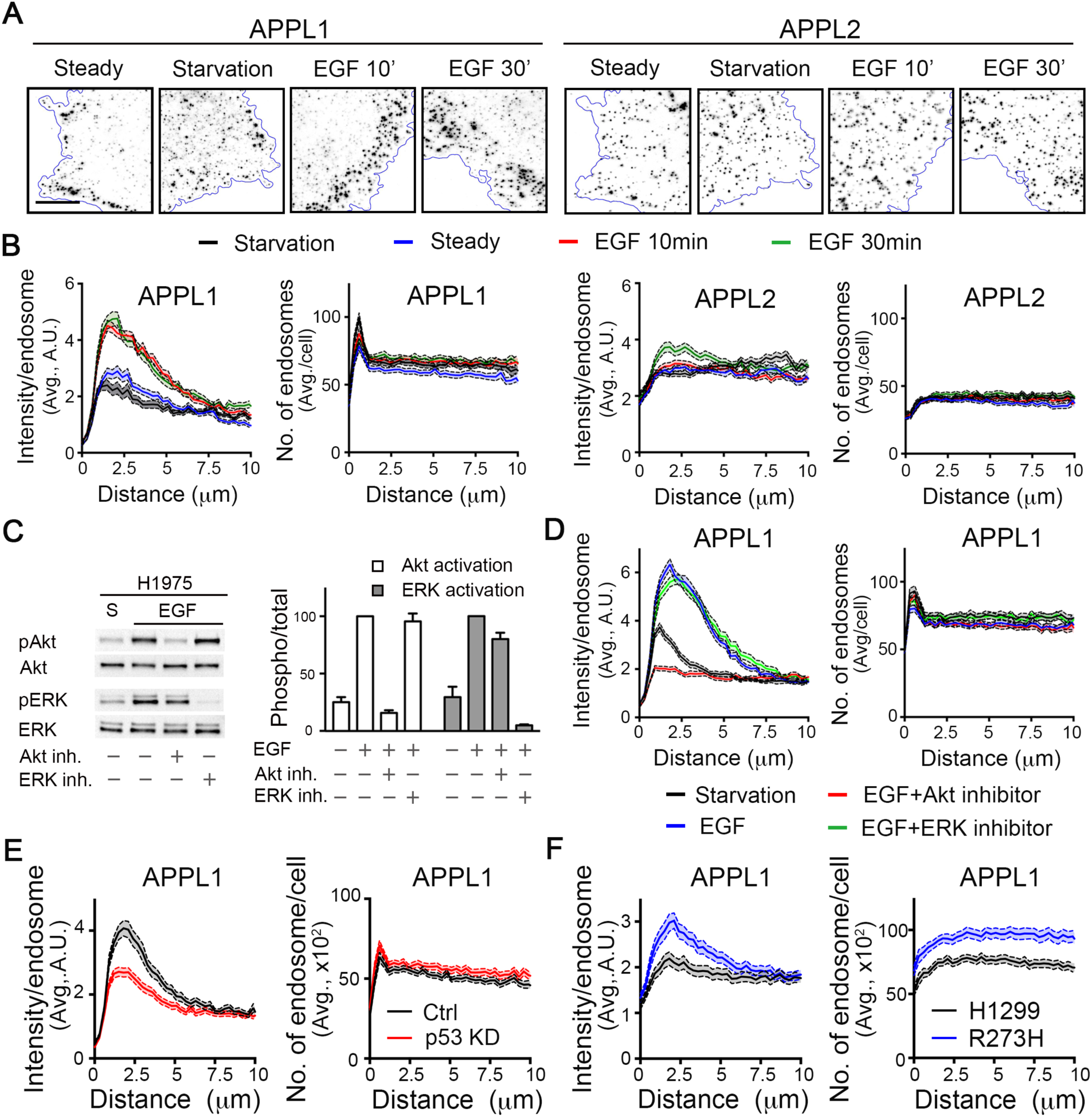
EGFR/Akt signaling regulates perimeter APPL1 endosomes in p53-driven cancer cells. (A) Representative ‘thick-TIRF’ immunofluorescence images of APPL1 or APPL2 endosomes in H1975 cells at steady state, or after starvation, and stimulation with EGF (20 ng/ml) for 10-or 30-minutes as indicated. Scale bar, 10μm. (B) Accompanying quantification of fluorescence intensity per endosome (left) and number (right) of APPL1-or APPL2-positive endosomes relative to their distance from the cell edge. Errors bars along each curve represent 95% confidence intervals. (C) Representative western blot and accompanying quantification (n=3) of phosphorylated Akt (Ser 473) and phospho-ERK in starved or EGF-stimulated treated with either Akt or ERK inhibitors. (D) Quantification of intensity/endosome or number of APPL1-positive endosomes in starved or EGF-treated H1975 cells, with or without 30-minute pre-treatment with 10 µM of either Akt (Akt Inhibitor X) or ERK (SCH772984) inhibitors. Quantification of intensity and number of APPL1-positive endosomes in control and p53 siRNA-treated H1975 cells (E) or parental and p53 R273H-expressing H1299 cells (F).

To accurately quantify the spatial distribution of APPL endosomes, we adapted our cmeAnalysis platform (Aguet et al., 2013) for high-throughput endosome detection and implemented distance analysis to assess endosome distribution relative to the cell edge (see Methods). Using these tools, we measured both the average intensity of individually detected endosomes and the total number of detected endosomes relative to the distance they reside from the cell edge (Figure 1B).

Starvation triggered a significant reduction in the fluorescent intensity of the subpopulation of APPL1 endosomes localized within ∼2.5 µm from the cell edge (hereafter referred to as ‘perimeter’ endosomes), without a corresponding change in the number of these peripherally-localized structures (Figure 1A, B). In contrast, the intensities of APPL2 endosomes did not change and they remained uniformly distributed throughout the cell periphery (Figure 1A, B). Stimulation with EGF resulted in a rapid (within 10 min) and dramatic accumulation of perimeter-localized, high-intensity APPL1 endosomes (Figure 1A). Interestingly, within cell clusters or in isolated migrating cells, the EGF-stimulated APPL1-positive perimeter endosomes appeared to accumulate towards the outside or leading edge (Supplemental Figure 1). The EGF-stimulated increase in perimeter APPL2 endosomes occurred at later time points (30 min) and to a much lesser extent (Figure 1B), indicating a specificity for the APPL1 isoform.

The observed EGF-dependent increase in the intensity of perimeter APPL1 endosomes was not accompanied by an increase in their numbers (Figure 1B). This suggests that EGF treatment either increased the expression of APPL1 and/or its recruitment to the membrane. We did not detect any EGF-dependent increase in total APPL1, but were able to detect EGF-dependent increases in the recruitment of APPL1 and its interactor Akt to cellular membranes (Supplemental Figure 2). Thus, we conclude that EGFR signaling regulates the selective recruitment of APPL1 to a subpopulation of perimeter-endosomes.

**Figure 2.**
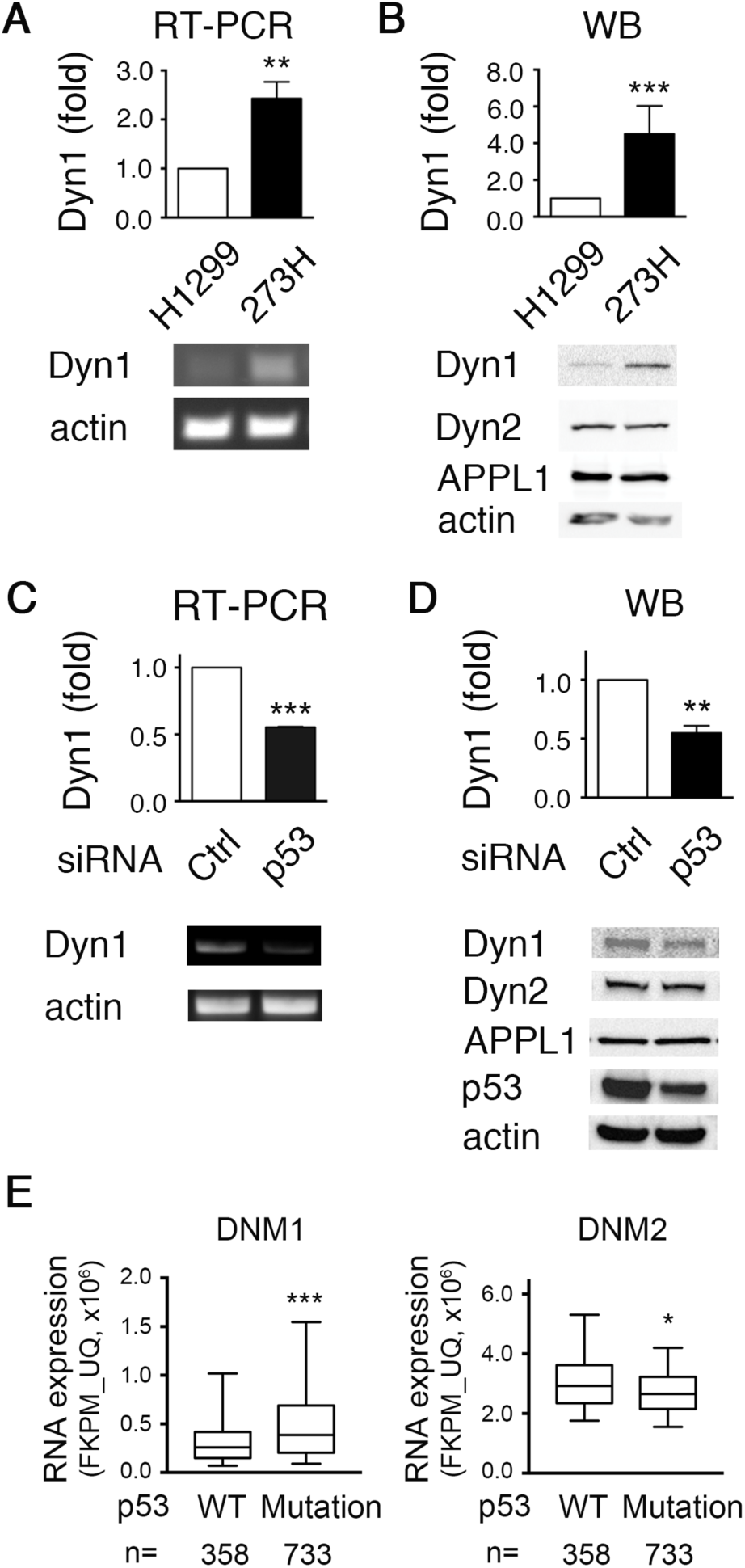
Dyn1 is upregulated downstream of mutant p53. (A) Quantification and representative RT-PCR of Dyn1 mRNA expression in parental and p53 R273H-expressing H1299 cells. (B) Quantification and representative western blots of Dyn1 expression in parental and p53 R273H H1299 cells. (C) Quantification and representative RT-PCR of Dyn1 mRNA in parental and p53 KD H1975 cells. (D) Quantification and representative western blots of Dyn1 expression in parental and p53 KD H1975 cells. All quantifications are from three independent experiments. Unpaired t-tests were used to assess statistical significance; **P<0.01, ***P<0.001. (E) Box plot of TCGA RNA expression profiles of DNM1 and DNM2 in human NSCLC tumors with wild-type (n=358) or any p53 mutation (i.e. deletions and/or mutations) (n=373). Mann–Whitney U-test was performed to compute significance; *P< 0.05, ***P< 0.001.

We next explored which of the two major signaling pathways downstream of EGFR, PI3K/Akt or Ras/MAPK/ERK, was required for the selective upregulation of perimeter APPL1 endosomes. To this end, cells were treated with either Akt inhibitor X or the ERK1/2 inhibitor SCH772984. Control experiments established that both compounds selectively inhibited their respective target kinases (Figure 1C). The EGF-stimulated accumulation of perimeter APPL1 endosomes selectively required Akt but not ERK1/2 activity (Figure 1D). Interestingly, even the residual subpopulation of perimeter APPL1 endosomes in starved cells were sensitive to Akt inhibition (Figure 1D), suggesting a direct link between Akt signaling and APPL1 recruitment. Taken together, our data establish that the accumulation of perimeter APPL1 endosomes in mutant p53-expressing H1975 cells is dependent on EGFR and/or Akt signaling.

Given that Akt is activated by a GOF mutant p53 downstream of EGFR (Muller et al., 2009), we examined the relationship between mutant p53 expression and perimeter APPL1 endosomes. Knockdown of mutant p53 in H1975 cells, which harbor homozygous p53 R273H mutations, resulted in a decrease in the intensity of perimeter APPL1 endosomes (Figure 1E). Correspondingly, ectopic expression of R273H mutant p53 in a p53-null NSCLC cell line, H1299, resulted in an increase in both the intensity and number of perimeter APPL1-positive endosomes (Figure 1F). These data link the accumulation of perimeter APPL1 endosomes to mutant p53 expression.

### Dyn1 is upregulated downstream of mutant p53

Intriguingly, the reported effects of mutant p53 expression on EGFR recycling (Mellman and Yarden, 2013; Muller et al., 2009; Muller et al., 2013) and our results showing a p53-dependent accumulation of APPL1 endosomes (Figure 1E,F), mirror those previously seen following activation of Dyn1 (Chen et al., 2017). These observations prompted us to investigate a possible role for Dyn1 in mediating p53-driven accumulation of perimeter APPL1 endosomes.

Strikingly, ectopic expression of GOF mutant p53 R273H in p53-null H1299 cells increased both mRNA (Figure 2A) and protein (Figure 2B) levels of Dyn1. siRNA-mediated knockdown of R273H p53 in H1975 cells resulted in a corresponding decrease in both Dyn1 mRNA (Figure 2C) and protein (Figure 2D) levels. These data suggest that Dyn1 might be a downstream target of p53. APPL1 protein levels were unaffected by mutant p53 expression (Figure 2B, 2D). Thus, as occurred upon EGF-stimulation, the p53-dependent increase in APPL1 endosomes likely reflects increased recruitment of APPL1 to nascent endosomal structures.

To connect these observations with human cancers, we mined NSCLC patient data in The Cancer Genome Atlas (TCGA) from Genome Data Commons (GDC) Data Portal and examined the relationship of Dyn1 mRNA expression and p53 mutation status. Downloaded patient data was split into two groups: patients with wild-type p53 and those with any p53 mutations. Consistent with our observations, the expression of Dyn1 mRNA in tumors with p53 mutations was significantly higher than in those expressing wild-type p53 (Figure 2E and Supplemental Table 1). In contrast, the expression of Dyn2 mRNA in tumors with p53 mutations was slightly reduced relative to those expressing wild-type p53. The above data suggest that mutant p53 expression alters early endocytic trafficking in cancer cells, in part, through transcriptional up-regulation of Dyn1, and consequently the accumulation of perimeter APPL1 endosomes.

### Dyn1 isoform-specific regulation of perimeter APPL1 endosomes

APPL1 endosomes have been reported to be regulated by CME (Sposini et al., 2017; Zoncu et al., 2009). Given the differential regulation of CME by dynamin isoforms (Reis et al., 2015; Srinivasan et al., 2018), we next tested their roles in regulating perimeter APPL1 endosome accumulation. Although siRNA-mediated knockdown of Dyn2 more potently inhibits CME in NSCLC cells than does Dyn1 knockdown (data not shown; but see Reis et al., 2015 and Srinivasan et al., 2018), only depletion of Dyn1 abrogated the accumulation of perimeter APPL1 endosomes in response to EGF signaling. Neither Dyn2 nor Dyn3 loss had any significant effect (Figure 3A and Supplemental Figure 3). The effect of Dyn1 depletion was specific to APPL1-endosomes as neither the distribution nor number of early EEA1-positive endosomes was affected (Supplemental Figure 4).

**Figure 3.**
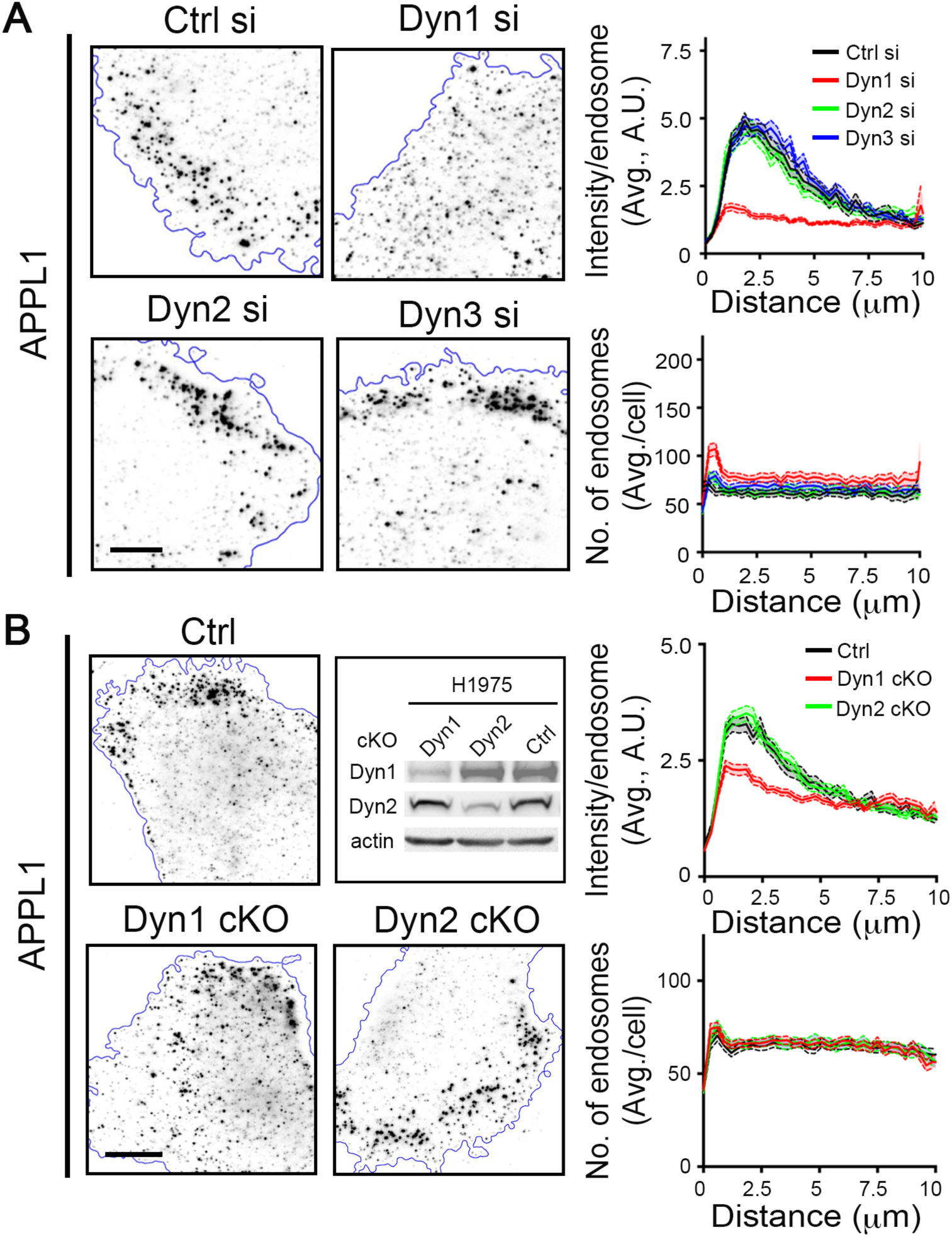
Dynamin-1 isoform-specific regulation of peripheral APPL1 endosomes. (A) Representative IF images and accompanying quantification of intensity and number of APPL1-positive endosomes in H1975 cells treated with scramble siRNA (Ctrl), or siRNA against Dyn1, Dyn2 or Dyn3. Cells were serum-starved and treated with EGF (20 ng/ml) for 10 min prior to fixation. (B) Representative IF images and accompanying quantification of intensity or number of APPL1-positive endosomes in control (cells infected with DD-Cas9 without guide sequence), DD-Cas9-Dyn1 (Dyn1 cKO) or DD-Cas9-Dyn2 (Dyn2 cKO) H1975 cells after four days of Shield-1 treatment. The representative western blot demonstrates Dyn1 and Dyn2 conditional knockout efficiency. Scale bar, 10μm.

**Figure 4.**
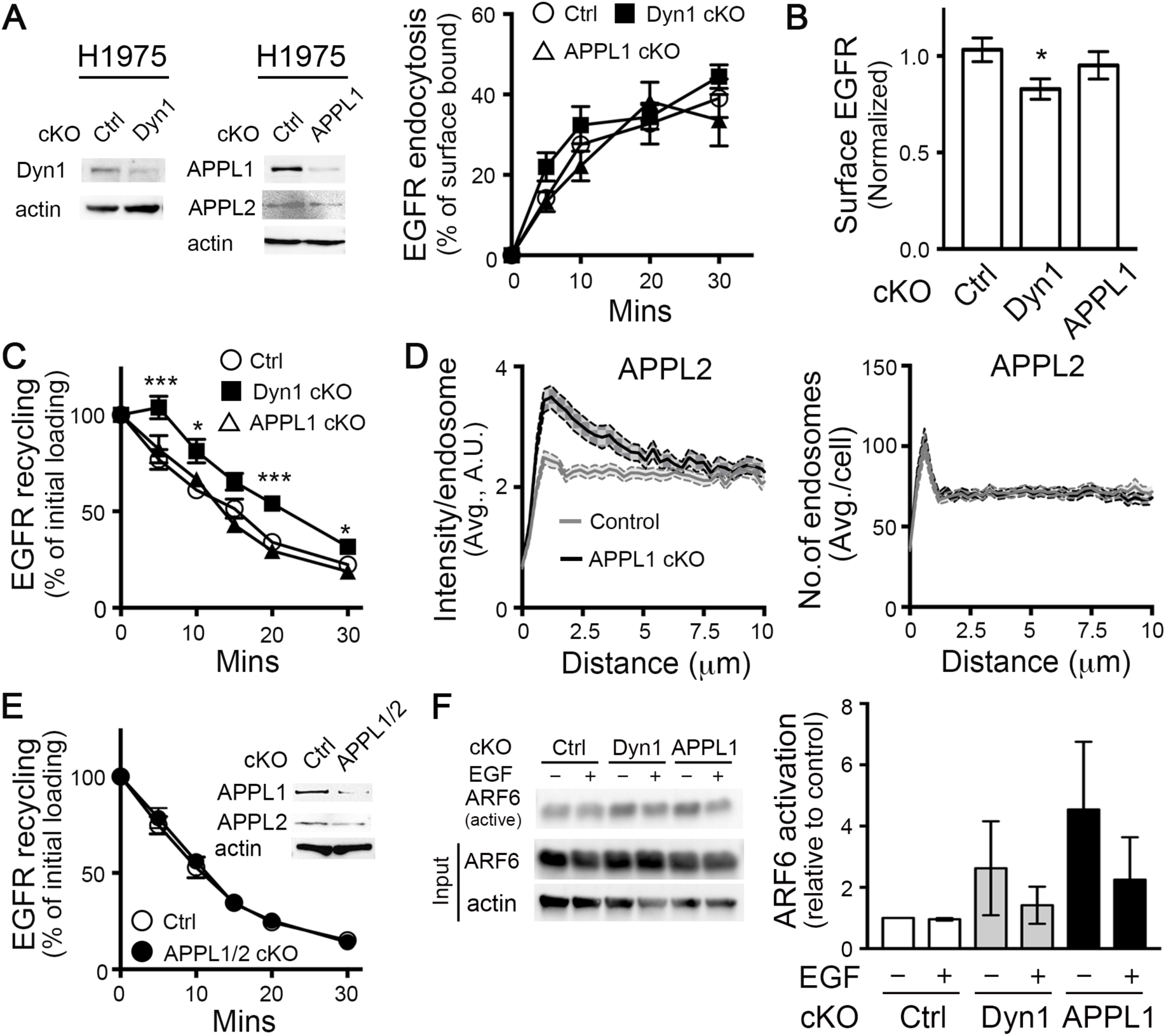
Dynamin-1 and APPL1 depletion reveal the plasticity of mechanisms regulating EGFR recycling. (A) Representative western blots showing efficiency of Dyn1 and APPL1 cKO and their effects on EGFR endocytosis. Percentage of internalized biotinylated-EGF was calculated relative to the initial surface bound. (B) Surface expression of EGFR in Control, Dyn1-or APPL1-cKO H1975 cells. (C) Effects of Dyn1 or APPL1 cKO on rapid recycling of EGFR. Cells were pulsed for 5 minutes with 20 ng/ml biotinylated-EGF, stripped, and reincubated at 37°C for the indicated times before measuring the remaining intracellular EGF. Percentage of recycled biotinylated-EGF was calculated relative to the initial loading. (D) Quantification of the intensity and distribution of APPL2 endosomes in APPL1 cKO H1975 cells. (E) EGFR recycling in Control, APPL1/2 double-cKO cells. Inset shows cKO efficiency. (F) Representative western blot and quantification showing compensatory upregulation of Arf6·GTP (i.e. active Arf6) in Dyn1 and APPL1 cKO cells. Error bars represent SEM (n=3). Unpaired t-tests were used to assess statistical significance; *P<0.05, **P<0.01, ***P<0.001.

To determine whether the role of Dyn1 in the accumulation of perimeter APPL1 endosomes was more general and/or solely dependent on mutant p53 expression, we evaluated the influence of depletion of each Dyn isoform on APPL1 and EEA1 endosomes in noncancerous and various cancer cell lines that express WT or mutant p53. These included two non-cancerous cell lines, ARPE-19 (human retinal pigment epithelial) and HBEC-3KT (human bronchial epithelial) that both express wild type p53, as well as a panel of cancerous cells, including NSCLC A549 cells that expresses wild type p53; MDA-MB-231 breast cancer cells that express homozygous p53 R280K; A375 and MV3 melanoma cells that both express wild type p53; and DU145 and PC3 prostate cancer cells that express heterozygous P223L, and V274F p53 and 138del p53, respectively. The number of APPL1 and EEA1 endosomes did not significantly change in any Dyn knockdown condition or cell line (data not shown). However, Dyn1 knockdown reduced the intensity of perimeter APPL1 endosomes in all cell lines tested, without affecting the distribution of EEA1 endosomes (Supplemental Figures 3-5). These data establish that the EGF-dependent accumulation of APPL1 perimeter endosomes selectively requires Dyn1 in both noncancerous and cancer cells. Thus, expression of GOF p53 mutations appear to co-opt this Dyn1/APPL1 endosome nexus.

Interestingly, neither Dyn2 nor Dyn3 knockdown inhibited the accumulation of perimeter APPL1 endosomes; rather, in some cells lines, Dyn2 and/or Dyn3 knockdown resulted in increased intensity of perimeter APPL1 endosomes (Supplemental Figures 3 and 5). It is possible that this reflects a compensatory upregulation of Dyn1, but this was not further investigated. Again, in all cases, EEA1 positive endosomes were unaffected (Supplemental Figure 4).

**Figure 5.**
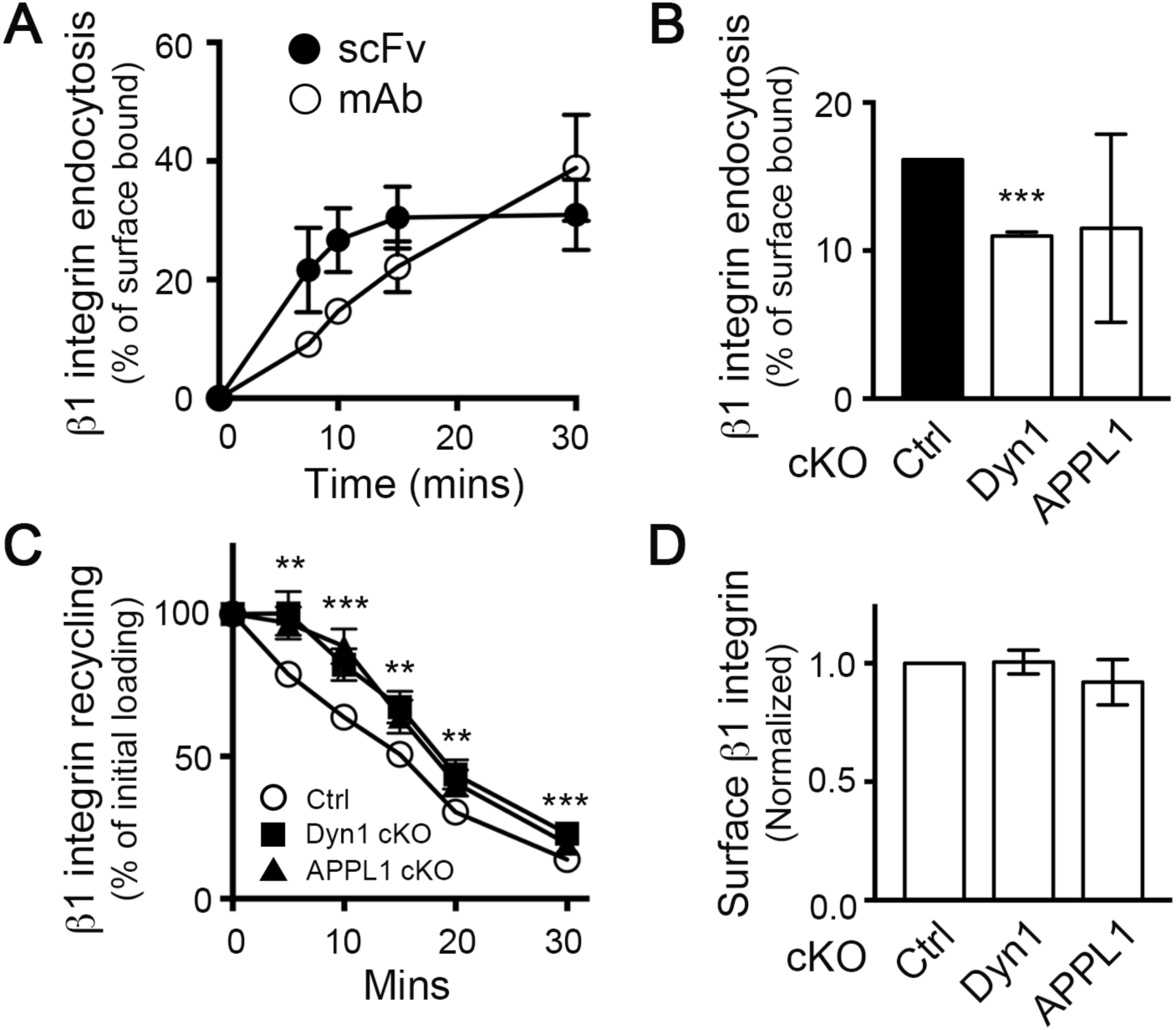
APPL1 and Dyn1 regulate β1 integrin recycling. (A) Endocytosis of biotinylated β1 integrin scFv (5μg/ml) or mAb K20 IgG in H1975 cells. Percentage of internalized antibody was calculated relative to the initial surface bound. (B) Endocytosis of biotinylated β1 integrin scFv (5μg/ml) was measured in control (Ctrl), Dyn1 cKO and APPL1 cKO H1975 cells. Shown is the percentage of internalized antibody after 10 minutes at 37°C. (C) Recycling of biotinylated β1 integrin scFv (5μg/ml) in Ctrl, Dyn1 cKO and APPL1 cKO H1975 cells. Percentage of recycled β1 integrin scFv was calculated relative to the initial loading (10 min). (D) Surface levels of β1 integrin in Dyn1 and APPL1 cKO cells normalized to Ctrl. All experiments represent n≥4. Error bars represent SEM. Unpaired t-tests were used to assess statistical significance; **P<0.01, ***P<0.001.

Given potential off-target effects of siRNA experiments, we confirmed these findings using inducible CRISPR (Clustered Regularly Interspaced Short Palindromic Repeats) knockout technology (Senturk et al., 2017) in H1975 cells. The advantage of inducible CRISPR over constitutive CRISPR is that it does not require clonal selection and expansion of knockout cells, which are susceptible to the induction/selection of compensatory mechanisms and clonal variation, especially problematic with heterogeneous cancer cell lines. We focused on Dyn1 and Dyn2, as Dyn3 is expressed at very low and frequently undetectable levels in most cell lines (Supplemental Figure 5) and is not required for APPL1 endosome distribution (Figure 3A and Supplemental Figure 3).

Consistent with siRNA experiments, conditional knockout (cKO) of Dyn1, but not Dyn2 suppressed perimeter APPL1-positive endosome accumulation compared to control H1975 cells infected with DD-Cas9 without targeting sequences (Figure 3B). Together, these results reveal a specific and isoform-selective role for the normally neuronally-enriched isoform, Dyn1, in regulating the accumulation of a mutant p53-and/or EGF-dependent spatially-restricted subpopulation of APPL1 signaling endosomes.

### Myo6 regulates the localization of perimeter APPL1 endosomes

Our data establish that Dyn1, which regulates APPL1 endosomes and early endocytic trafficking when activated in cancer cells, is upregulated upon expression of mutant p53. We next sought additional steps in the mechanism that might contribute to the mutant p53-driven effects on APPL1 endosomes. A recent study showed that Myo6, a minus end-directed actin motor, is required for the peripheral localization of APPL1 endosomes and their role in Akt activation (Masters et al., 2017). Interestingly, Myo6 has also been reported as a transcriptional target of wild-type and the R294S GOF mutant p53 (Jung et al., 2006). We extended these results showing that depletion of the R273H p53 mutant by siRNA in H1975 cells also decreased Myo6 expression (Supplemental Figure 6A). Importantly, siRNA-mediated depletion of Myo6 suppressed perimeter APPL1 endosome accumulation (Supplemental Figure 6B), and correspondingly attenuated EGF-stimulated Akt signaling in H1975 cells (Supplemental Figure 6C). Together, these data suggest that Myo6 expression is an additional mechanism downstream of mutant p53, required to modulate the localization of perimeter APPL1 endosomes and Akt signaling activity.

**Figure 6.**
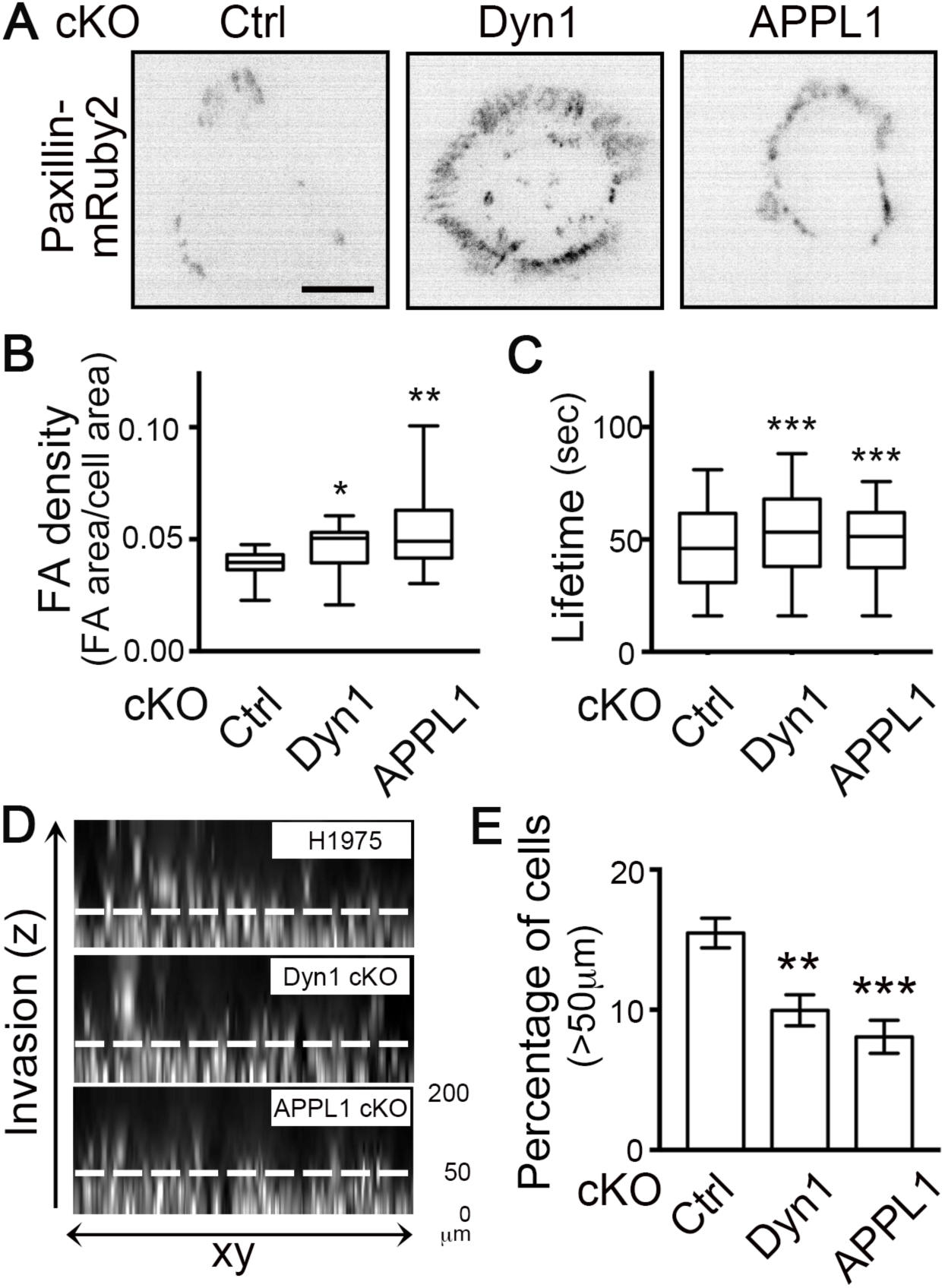
Dyn1 and APPL1 modulate focal adhesion turnover, cell migration and invasion. (A) Representative images of H1975 cells expressing Paxillin-mRuby2 in Ctrl, Dyn1 and APPL1 cKO conditions. Scale bar, 10μm. (B) Focal adhesion density in control (Ctrl), Dyn1 and APPL1 cKO H1975 cells. Box and whisker plots represent the 5-95^th^ percentile and indicated mean and outliers of the data from three independent experiments (n>49 images with more than one cell per image). (C) Focal adhesion lifetime in control (Ctrl), Dyn1 and APPL1 cKO H1975 cells. Data from three independent experiments (n>11 movies). Mann–Whitney U-test was performed to compute significance: *P<0.05; **P<0.01; ***P<0.001. (D) Representative Hoescht-stained images and (E) Quantification of migration and invasion of Ctrl, Dyn1 cKO, APPL1 cKO H1975 cells evaluated in an inverted three-dimensional collagen matrix. Values are mean ± SEM, n=3. Statistical significance was analyzed by Student’s t test; **P< 0.01, ***P<0.001.

### Dynamin1 and APPL1 depletion reveal the plasticity of mechanisms regulating EGFR recycling

We previously identified a positive feedback loop involving Akt and Dyn1 activation that regulates EGFR trafficking and signaling (Chen et al., 2017; Reis et al., 2015). Others have recently shown a role for APPL1 endosomes in regulating GPCR recycling (Sposini et al., 2017). Therefore, we next tested for the function of Dyn1-dependent APPL1 perimeter endosomes in EGFR trafficking. Conditional depletion of neither Dyn1 nor APPL1 by DD-Cas9 altered the rate of EGFR endocytosis (Figure 4A). However, cKO of Dyn1, but not APPL1, induced a slight reduction of surface EGFR levels (Figure 4B), suggesting a defect in receptor recycling.

To directly measure the potential short-circuited, rapid recycling of EGFR from perimeter endosomes, we developed a sensitive assay to follow the fate of a brief, 5-minute pulse of internalized EGF (see STAR Methods). As expected from the observed decrease in surface EGFR, cKO of Dyn1 resulted in an ∼5 min lag before we could detect EGFR recycling (Figure 4C), suggesting progression of the EGF pulse deeper along the endosomal pathway prior to returning to the cell surface.

Given that APPL1 endosome distribution is dependent on Dyn1, we were surprised that APPL1 cKO did not alter the kinetics of EGFR recycling (Figure 4C). Therefore, we looked for compensatory mechanisms that might account for the unexpected differential effects of Dyn1-and APPL1-knockout on EGFR recycling. Indeed, we found that APPL2 accumulated on perimeter endosomes upon APPL1 depletion (compare Figures 1B and 4D). Depletion of APPL1 did not alter the number of APPL2 endosomes or APPL2 protein levels (Figures 4A, D). These data suggested that APPL2 might be recruited to perimeter endosomes in compensation for the loss of APPL1. To test this, we generated double APPL1/APPL2 DD-Cas9 H1975 cells and found, surprisingly, that even simultaneous depletion of both APPL isoforms did not affect EGFR recycling (Figure 4E). We conclude that either Dyn1 and APPLs have differential effects on EGFR recycling, or that yet other compensatory mechanisms exist.

EGFR recycling can be regulated by the small GTPase Arf6 (Allaire et al., 2013). To determine whether changes in Arf6 activity might constitute another mechanism of compensation, we measured Arf6 activity in control and APPL1 cKO cells using an Arf6·GTP pull-down assay (Cohen and Donaldson, 2010). Interestingly, steady-state Arf6 activity was elevated in both Dyn1-and APPL1-knockout cells when compared to control (Figure 4F), although EGF treatment decreased Arf6 activity in both cKO cell lines. Together these data suggest a high degree of plasticity in early endocytic trafficking and recycling of activated EGFR, as has also been observed for its uptake via CME (Goh et al., 2010). Nonetheless, that these compensatory mechanisms are induced in response to depletion of Dyn1 and/or APPL1 suggests a role for both Dyn1 and perimeter APPL1 endosomes in regulating the rapid recycling of EGFR.

### Dyn1 and APPL1 are required for β1 integrin recycling

Given the complexity of early endocytic trafficking and sorting of the EGFR, especially in cancer cells (Tomas et al., 2014), we next studied β1 integrin trafficking, as these adhesion receptors are critical for p53-dependent changes in cancer cell migration and invasion (Muller et al., 2014; Muller et al., 2013). However, to accurately measure rapid, early recycling of β1 integrins required the development of more sensitive assays than those previously used. For example, in studies using surface biotinylation, cells were incubated for ≥30 min to accumulate sufficient internalized signal for subsequent recycling measurements (Arjonen et al., 2012; Caswell et al., 2008; Muller et al., 2009; Muller et al., 2014; Muller et al., 2013; Roberts et al., 2001). Other studies used surface bound mAbs to follow β1 integrin trafficking, which can inhibit rapid recycling due to clustering of the bivalent Ab-bound receptor (Weissman et al., 1986). To circumvent these problems, we developed a single chain antibody variable fragment (scFv) derived from mAb K20, a well-characterized, non-function perturbing, monoclonal anti-β1 integrin antibody (Amiot et al., 1986; Byron et al., 2009; Takada and Puzon, 1993) that can be used to measure β1 integrin uptake and recycling (Lakoduk et al, ms in preparation). A comparison of internalization rates of the β1 integrin scFv with its parent mAb reveals that uptake of mAb K20 continues linearly for ≥30 min, whereas the scFv is internalized faster and reaches equilibrium after 15 minutes (Figure 5A). These data are consistent with rapid and efficient recycling of scFv.

In contrast to EGFR, conditional depletion of Dyn1 inhibited β1 integrin internalization, as did APPL1 depletion, although to a more variable extent (Figure 5B). A cargo-selective role for Dyn1 in CME has been previously reported (Reis et al., 2017). More importantly, the rapid recycling of β1 integrins, which we could measure directly after a 10-min internalization pulse of scFv, was equally inhibited following either Dyn1 or APPL1 depletion (Figure 5C), both of which resulted in a 5 min lag before recycling could be detected. In combination, these two effects resulted in unchanged levels of surface β1 integrin (Figure 5D). Together these data indicate that both Dyn1 and APPL1 endosomes are required for rapid β1 integrin recycling.

### Dyn1 and APPL1 modulate focal adhesion turnover, migration and invasion

We have shown that Dyn1 is upregulated downstream of mutant p53 and is required for perimeter APPL1 endosome accumulation downstream of EGFR/Akt signaling to increase fast recycling of EGFR and β1 integrin. The dynamic turnover of cell surface integrins regulates focal adhesions (FAs) that form at the edges of cells and are critical for cell migration (Figure 6A). Therefore, to link these observations to the p53-dependent changes in cancer cell invasion, we first used paxillin immunofluorescence in fixed cells to assess the density of FAs (FA area/cell area) at steady-state in mutant p53-expressing H1975 cells. cKO of Dyn1 and APPL1 resulted in increased FA density compared to control (Figure 6B). Interestingly, cKO of Dyn1 and APPL1 resulted in increased adhesion density at cell edges (Figure 7A, B), from where perimeter APPL endosomes are lost. Depletion of Dyn1 and APPL1 also resulted in significantly increased FA lifetimes, which was measured by expressing low levels of mRuby2-tagged paxillin (Figure 6C). Together, our results demonstrate a role for Dyn1 and perimeter APPL1 endosomes in regulating integrin and adhesion turnover in cancer cells.

**Figure 7.**
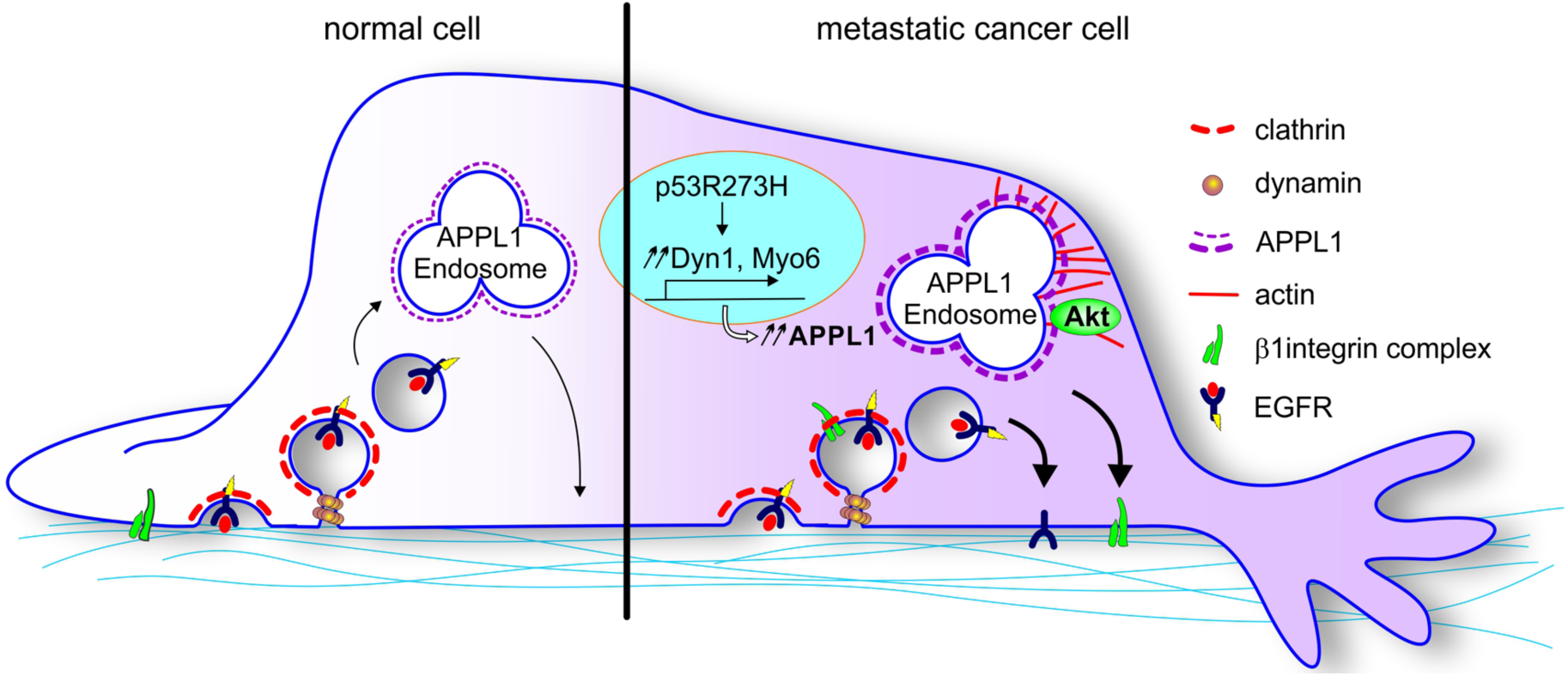
A Dynamin-1 and APPL1 endosome nexus regulates β1 integrin recycling downstream of EGFR, Akt and mutant p53. Dyn1 and Myo6, whose expression levels are increased in cells expressing mutant p53, regulate the accumulation of APPL1 ‘signaling’ endosomes at the cell perimeter to accelerate EGFR and β1 integrin recycling. A positive-feedback loop involving APPL1-dependent Akt signaling and activation of Dyn1 is established. Together, these cancer cell-induced changes in endocytic membrane trafficking lead to increased migration and invasion.

Finally, we investigated the effect of depletion of Dyn1 and APPL1 on cell migration/invasion using an inverted invasion assay in which cells migrate upwards towards EGF through a collagen/fibronectin layer. Conditional KO of either Dyn1 or APPL1 significantly inhibited cell invasion beyond 50 µm (Figure 6D, E). Together, our data establish that Dyn1-regulated perimeter APPL1 endosomes modulate cancer cell migration and invasion through alterations in early endocytic trafficking of EGFR and β1 integrin.

## Discussion

Solid tumors, initially triggered by oncogenic transformation, require numerous adaptations to progress to the aggressive metastatic state. Among these are alterations in receptor signaling and endocytic membrane trafficking that can lead to enhanced proliferation, survival, and invasive properties of the evolved cancer cell (Lanzetti and Di Fiore, 2017; Schmid, 2017). Previous studies have shown that p53 mutations linked to numerous cancers can alter endocytic trafficking of RTKs and integrins resulting in increased cell migration and metastasis (Muller et al., 2009; Muller et al., 2013). Here, we identify molecular mechanisms underlying these effects. Our data establishes that Dyn1 and Myo6 upregulation downstream of gain-of-function p53 mutations results in the accumulation of a spatially-restricted subpopulation of APPL1-positive endosomes and increased Akt signaling, which in turn activates Dyn1. Together this positive feedback loop and the Dyn1/perimeter APPL1 endosome nexus culminates in increased migration and invasion in cancer cells. Our findings exemplify how crosstalk between receptor signaling and endocytic trafficking, as well as selective activation of endocytic protein isoforms, can establish feedback loops that contribute to aggressive cancer cell behaviors.

We have previously reported that APPL1 endosome-dependent Akt/GSK3β signaling activates normally quiescent Dyn1 in non-neuronal cells to increase rates of clathrin coated pit initiation and maturation (Chen et al., 2017; Reis et al., 2015; Srinivasan et al., 2018). Here we show that both Akt activity and Dyn1 are also required for the accumulation of perimeter APPL1 endosomes. Together, these findings establish that mutant p53 triggers a positive feedback loop, through upregulation of Dyn1 and Myo6, accumulation of perimeter APPL1 endosomes and concomitant increased Akt activity, that alters early endocytic trafficking, signaling and cell migration (Figure 7).

Recent studies have shown that APPL1 endosomes function in the rapid recycling of a subset of GPCRs (Jean-Alphonse et al., 2014; Sposini et al., 2017) and that this recycling pathway is inhibited by a negative feedback loop driven by cAMP-dependent activation of protein kinase A. Together with our findings, these results suggest that APPL1 endosomes are central players in the crosstalk between specific signaling receptors and their endocytic trafficking. That spatially-restricted subpopulations of APPL1 endosomes are differentially regulated adds another layer of complexity to this crosstalk. The assays we have developed to measure rapid recycling and peripheral endosome distribution should help to unravel this complexity.

Altered β1 integrin recycling is associated with promoting cancer cell dissemination (Jones et al., 2006; Muller et al., 2009; Paul et al., 2015). Canonically, β1 integrin can recycle back to the plasma membrane through either a short-loop (fast) or a perinuclear long-loop (slow) pathway (Caswell et al., 2009; Ivaska and Heino, 2011; Paul et al., 2015). The former is selectively regulated by growth factor receptor signaling (Arjonen et al., 2012; Fang et al., 2010; Onodera et al., 2012), although the mechanisms for this regulation have not been understood. Previous studies have shown the polarized redistribution of RCP-positive recycling endosomes at leading extensions of cells migrating in 3D microenvironments towards a chemotactic signal (Caswell et al., 2008). Thus, it is possible that the recruitment of APPL1 endosomes to the cell edge we observe in response to global EGF stimulation might be more polarized in response to directional chemotactic signals. Indeed, we observe a polarized distribution towards the outer edges of cell clusters (Supplemental Figure 1). Together, our findings support a molecular mechanism for the necessary role of growth factors or serum in regulating a spatially-restricted subpopulation of endosomes involved in the fast recycling of β1 integrin.

APPL1 was initially identified as an Akt interacting protein (Mitsuuchi et al., 1999) involved in cell survival and proliferation (Ding et al., 2016; Schenck et al., 2008). However, the role of APPL1 in regulating cell migration is still controversial. Recent studies suggest a negative role of APPL1 in cell migration and Akt signaling in Ras-driven HT1080 fibrosarcoma cells (Broussard et al., 2012; Diggins et al., 2018). In contrast, other reports support a role for APPL1 in increasing cell migration in leptin-stimulated cancer cells (Ding et al., 2016) and in HGF-stimulated mouse embryo fibroblasts (Tan et al., 2016). Our results support a positive role for a specific subpopulation of APPL1 endosomes in cell migration in EGF-stimulated, mutant p53-driven H1975 cells. Differences in cell types used and the signaling pathways studied may account for these different findings. More importantly, these differences reflect the heterogeneity and plasticity in mechanisms driving cancer cell progression.

Cancer cells exhibit significant heterogeneity and there are no doubt multiple pathways by which they can adapt their early endocytic trafficking during tumor progression. Indeed, the plasticity of the mechanisms governing early endocytic trafficking in cancer cells was evident in the induction of compensatory mechanisms governing EGFR recycling upon conditional CRISPR-mediated knockout of Dyn1 or APPL1 (e.g. ARF6 activation in Dyn1 and APPL1 cKO cells, and APPL2 redistribution in APPL1 cKO cells). These observations suggest a flexibility of endocytic and signaling systems that can lead to increased plasticity and adaptation in cancer cells.

Both factors–the induction of compensatory mechanisms and heterogeneity–can complicate mechanistic analyses of cancer cells. State-of-the-art CRISPR-based DNA editing has greatly advanced our understanding of gene functions and improves off-target issues associated with siRNA experiments (Hsu et al., 2014; Setodji et al., 2017). However, most constitutive CRISPR knockout studies require single cell clonal expansion and are therefore susceptible to inherent problems of clonal variation and compensation arising from prolonged cell culture. We adopted a conditional CRISPR knockout system to circumvent these issues. However, and unexpectedly, we still observed that in just four days, cancer cells can adapt quickly to manipulation.

The extent of our current knowledge of the molecular mechanisms governing early endocytic trafficking provides a foundation for identifying and testing possible compensatory mechanisms. As we have demonstrated, applying this growing knowledge base regarding membrane trafficking and its regulation will improve our understanding of cancer cell plasticity and heterogeneity. Reciprocally, our studies on the regulation of early endocytic trafficking in cancer cells have revealed added complexity with regard to the spatial organization and functional diversity of early endosomes.

Together our findings suggest that tunable crosstalk between endosomal recycling and receptor signaling networks, in combination with feedback loops, can be activated in cancer cells to enhance their metastatic traits. That the components of the endocytic machinery that regulate APPL1 signaling endosomes and rapid recycling are targeted by gain-of-function p53 mutations frequently associated with cancer speaks to the evolutionary advantage of manipulating this pathway. More generally, our data provide mechanistic insight into how selective activation of endocytic isoforms can alter endosomal recycling and receptor signaling to promote the adaptation required for aggressive phenotypes in cancers.

## Author contributions

A.M.L. and P.-H.C. designed the project and performed the experiments. A.M.L, P.-H.C. and S.L.S. discussed and analyzed the results. P.R. generated new computational analysis software for endosome IF experiments. M.M. assisted with experiments and analyses. H.M.G. assisted with virus preparation and cell line generation. A.M.L., P.-H.C. and S.L.S. wrote the manuscript with input from all authors.

## Acknowledgments

We are grateful to Dr. Szu-Chin Fu (Department of Pharmacology, UTSW Medical Center) for help in downloading, analysis and discussion of TCGA data from GDC, and Dr. Bo-Jui Chang (Department of Cell Biology, UTSW Medical Center) for image analysis of inverted invasion assay. We also appreciate Dr. Sangyoon J. Han (Department of Biomedical Engineering, Michigan Technological University) and Dr. Andrew Jamieson (Department of Bioinformatics, UTSW Medical Center) for helpful discussion regarding the focal adhesion analysis software, and all additional help from Schmid lab members. This research was supported by NIH grants GM73165 to S.L.S and Gaudenz Danuser and GM42455 to S.L.S. P.-H.C. was partly supported by Taiwan National Science Council Grant 103-2917-I-564-029.

## Methods

Detailed methods are provided in the online version of this paper and include the following:

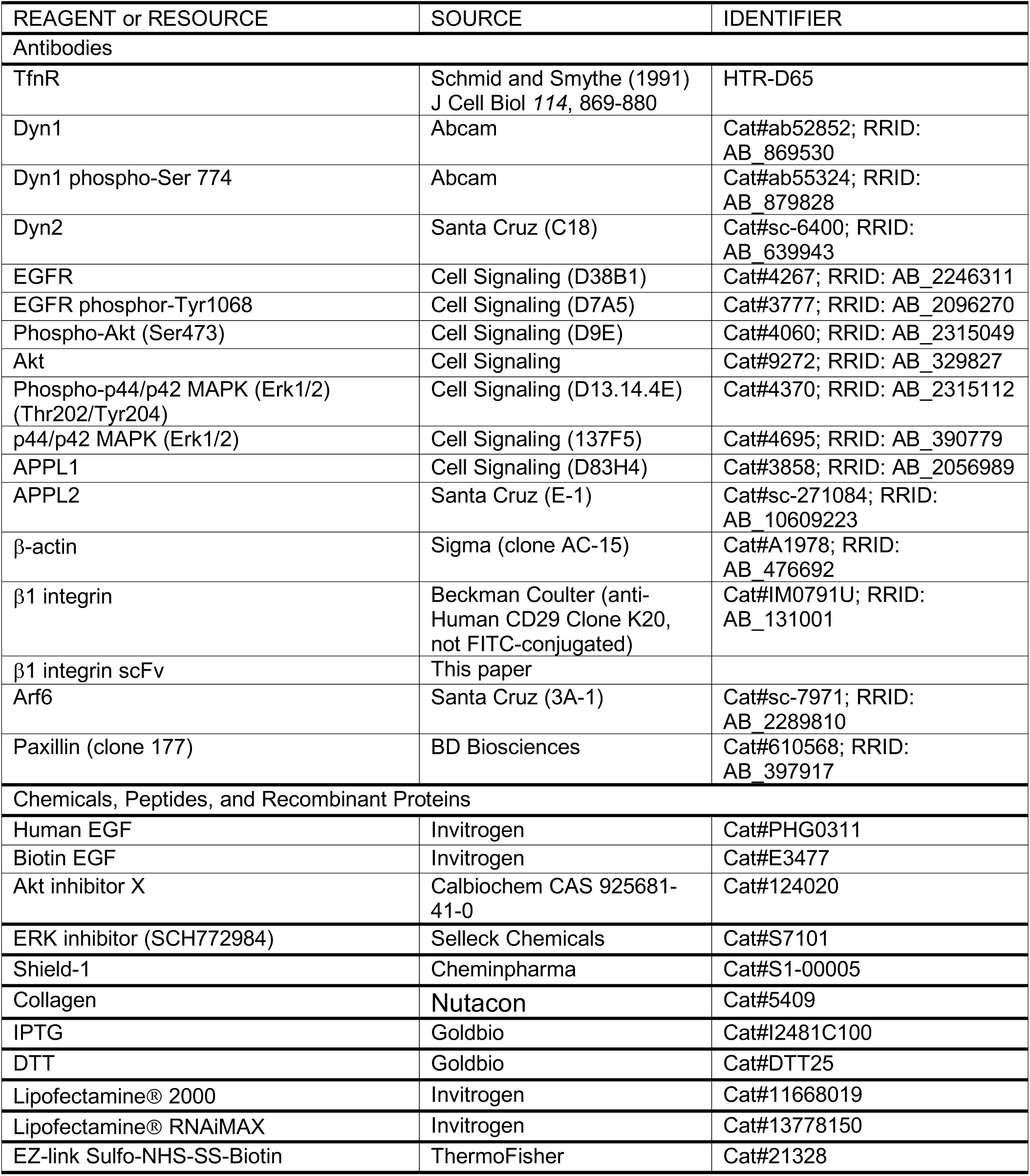
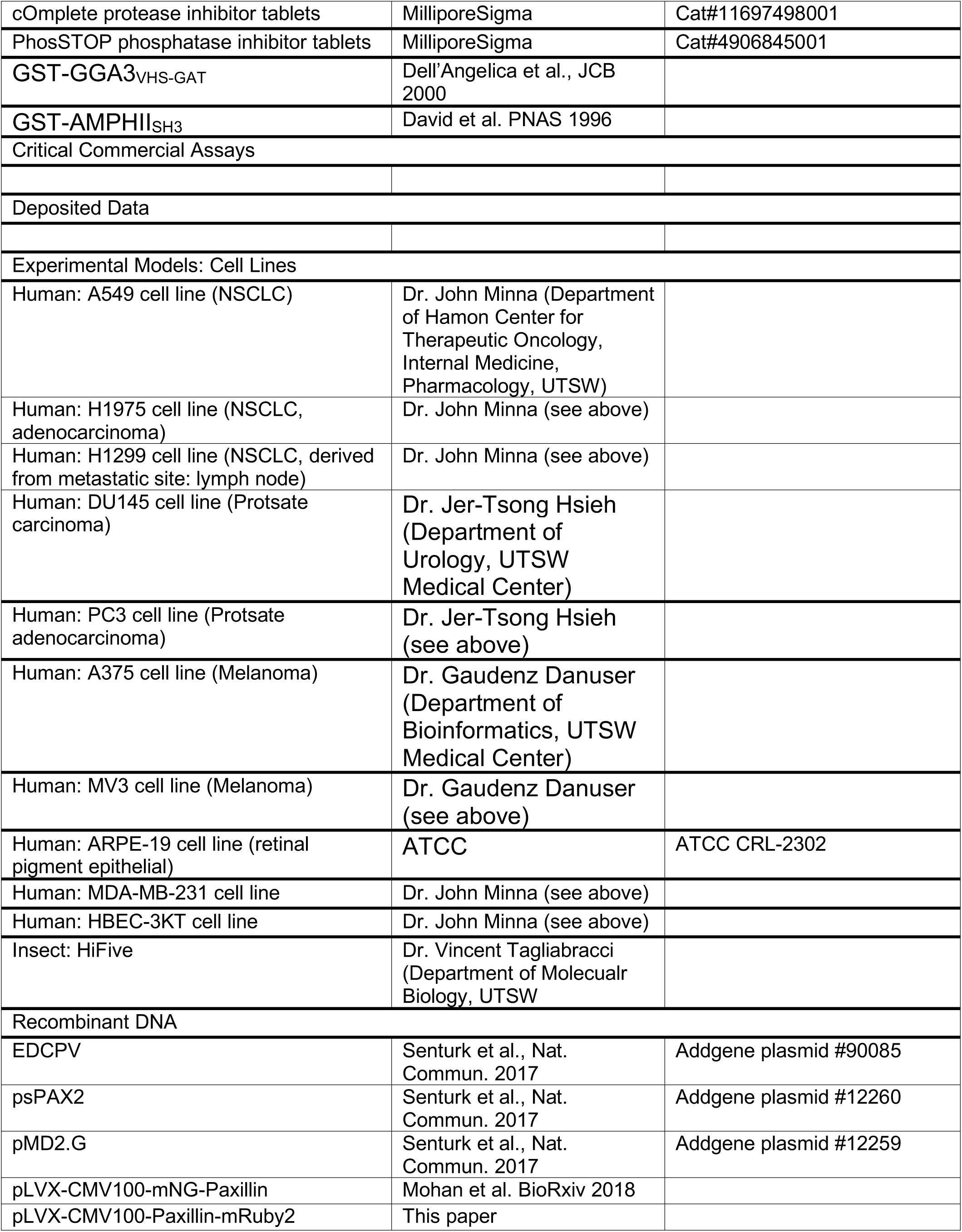
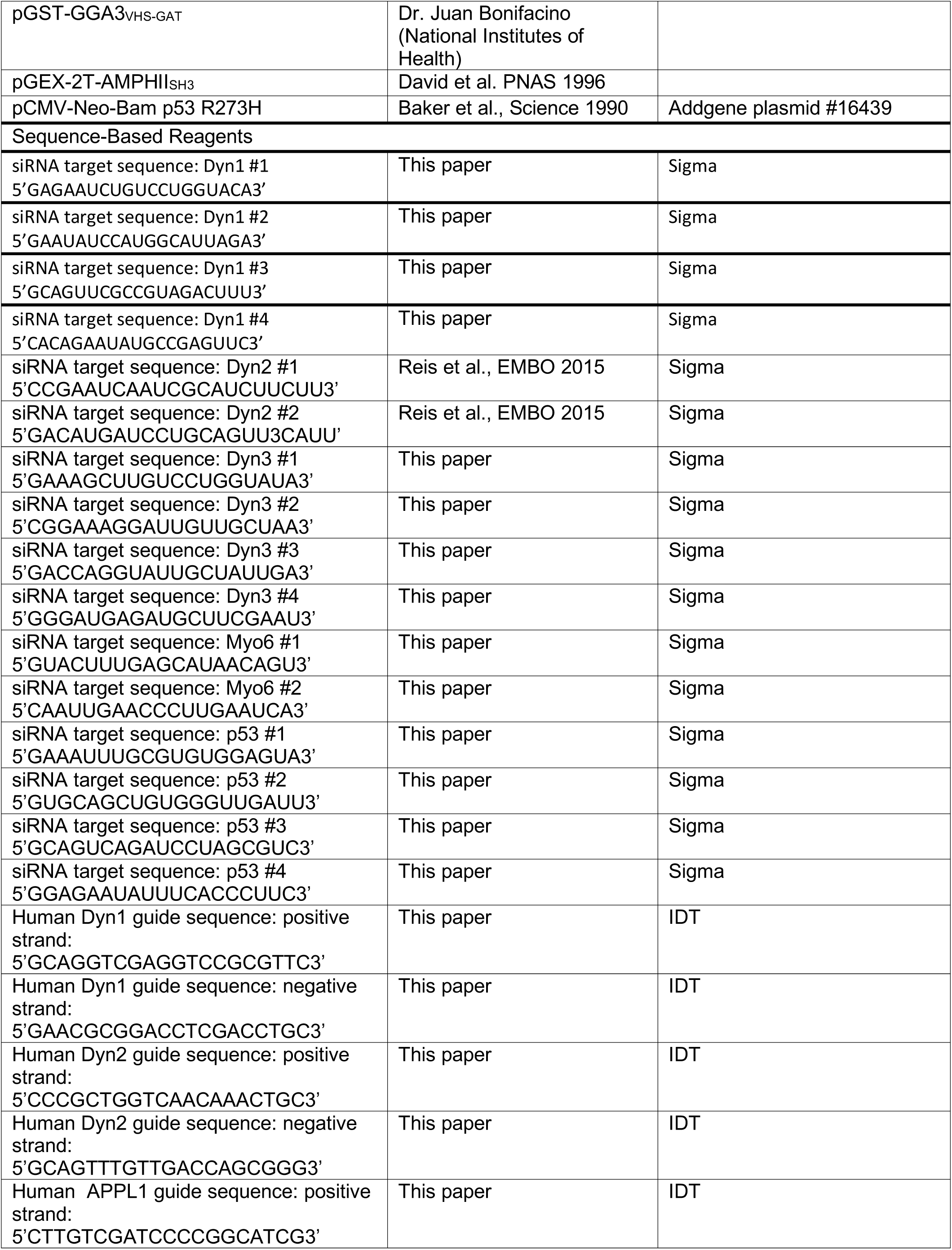
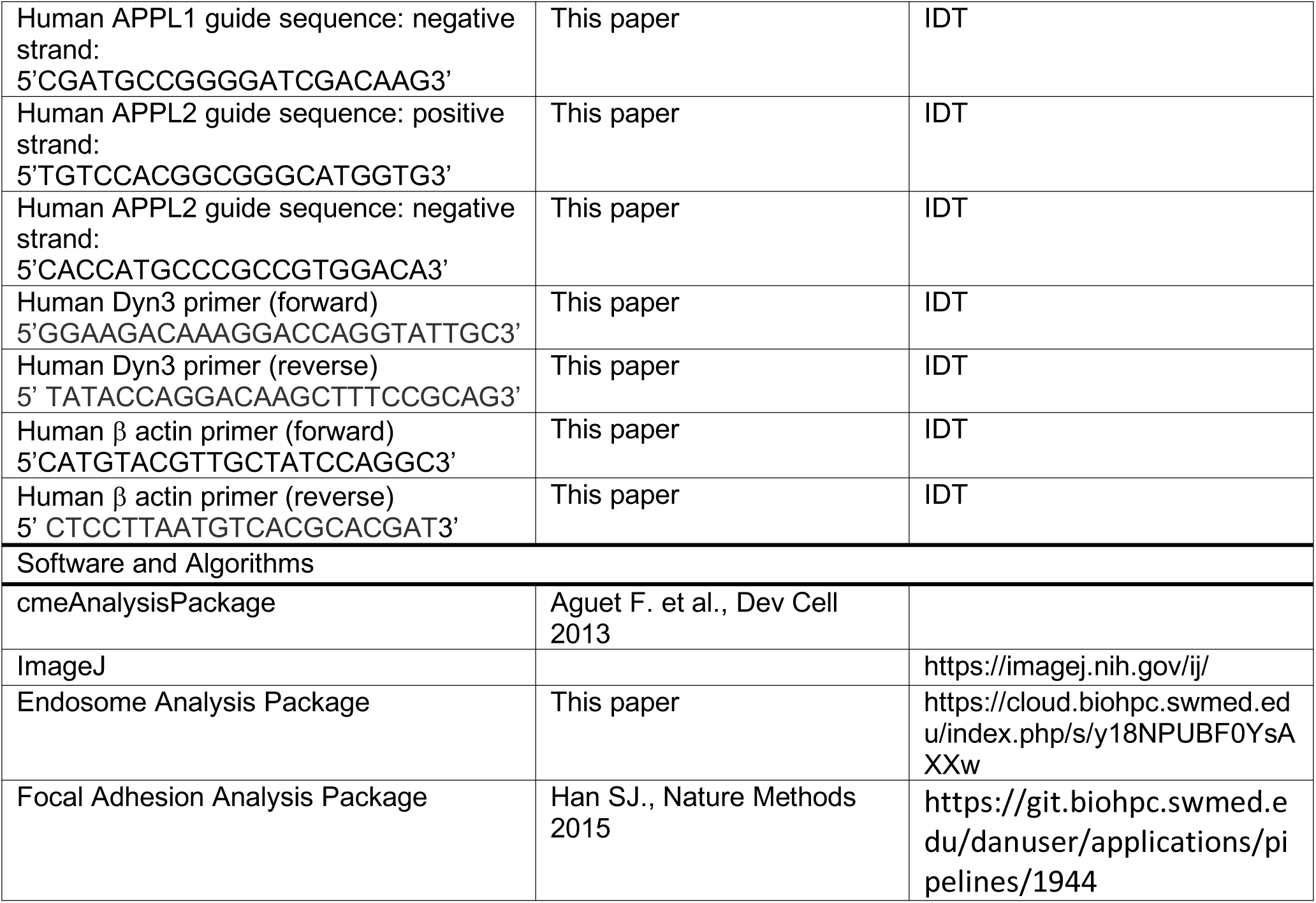
KEY RESOURCES TABLE.

## METHOD DETAILS

### Cell culture

H1299, H1975, and A549 cells were kindly provided by Dr. John Minna (Department of Hamon Center for Therapeutic Oncology, Internal Medicine, Pharmacology, UTSW Medical Center). DU145 and PC3 cells were kindly provided Dr. Jer-Tsong Hsieh (Department of Urology, UTSW Medical Center). A375 and MV3 cells were kindly provided by Dr. Gaudenz Danuser (Department of Bioinformatics, UTSW Medical Center). A549, DU145 and PC3 cells were maintained RPMI 1640 medium (Gibco) supplemented with 10% of FBS. HBEC-3KT cells were maintained Keratinocyte-SFM medium (ThermoFisher) supplemented with bovine pituitary extract (ThermoFisher). ARPE-19 cells were maintained in F12/DMEM medium (ThermoFisher) supplemented with 10% of FBS. MDA-MB-231, A375 and MV3 cells were maintained in DMEM medium (ThermoFisher) supplemented with 10% of FBS. H1975 and derived cell lines were maintained in biotin-free RPMI 1640 medium (USBiological) to reduce biotin background for biochemical assays, supplemented with 10% of FBS

### Cell engineering

p53 R273H reconstituted H1299 cells were generated by transfecting with pCMV-Neo-Bam p53 R273H plasmids and selected in completed medium containing G418 (1mg/ml, ThermoFisher).

Isogenic inducible H1975 derived DD-Cas9-Dyn1, DD-Cas9-Dyn2, DD-Cas9-APPL1 and DD-Cas9-APPL2 cells were generated by expansion of Venous positive cells following FACS sorting of cells expressing EDCPV (Addgene plasmid #90085, Senturk et al., 2017) harboring human Dyn1 guide sequence (Positive strand,5’GCAGGTCGAGGTCCGCGTTC3’;Negativestrand, 5’GAACGCGGACCTCGACCTGC3’), human Dyn2 guide sequence (Positive strand,5’CCCGCTGGTCAACAAACTGC3’;Negativestrand, 5’GCAGTTTGTTGACCAGCGGG3’), human APPL1 guide sequence (Positive strand,5’CTTGTCGATCCCCGGCATCG3’;Negativestrand, 5’CGATGCCGGGGATCGACAAG3’) or human APPL2 guide sequence (Positive strand,5’TGTCCACGGCGGGCATGGTG3’;Negativestrand, 5’CACCATGCCCGCCGTGGACA3’). Control cells expressed the DD-Cas9 construct without a guide sequence.

### Immunofluorescence analysis

Briefly, 22×22 mm glass slips were coated with 0.01% poly-L-lysine for 10 minutes at RT, washed with PBS, then coated with 0.02% gelatin (in PBS containing 2% sucrose) at 37°C for 30 minutes, followed by crosslinking with 1% paraformaldehyde for 15 minutes at RT. Gelatin-coating slips were extensively washed with PBS and then incubated in complete medium at 4°C for overnight. 3×10^5^ cells were seeded on gelatin-coated slips overnight, then washed with PBS, and starved in serum-free medium for 1 hour. After that, 20ng/ml of EGF was added to cells for 10-30 minutes, as indicated. Cells were washed once in PBS, fixed in 4% paraformaldehyde for 30 minutes at 37°C, quenched in 100mM glycine for 5 minutes, washed with PBS, permeabilized in 0.2% Triton X-100 for 10 minutes at RT and then blocked in 5% BSA/PBS for 1 hour at RT. The fixed and permeabilized cells were then incubated with primary antibodies (1:250 dilution in Q-PBS which contains 0.2% BSA, 0.001% saponin and 0.01% glycine) at 4°C overnight. After three PBS washes, the cells were incubated with Alexa Fluor-conjugated secondary antibodies (1:1000 dilution in Q-PBS) at 37°C for 1 hour. After three PBS washes, the cells were mounted in PBS and imaged by TIR-FM.

### siRNA and Plasmids transfection

Plasmids and siRNA were transfected according to the manufacturer’s recommendations using Lipofectamine 2000 and Lipofectamine RNAiMAX respectively (Invitrogen). Briefly, 110 pmol of the indicated siRNA and 6.5 μl of Lipofectamine RNAiMAX reagent were diluted in 100 μl of OptiMEM medium (ThermoFisher). Then, the diluted siRNA was added to diluted Lipofectamine RNAiMAX reagent and incubated for 5 minutes at room temperature. The mixed siRNA-lipid complex was added to each well of a 6-well plate containing cells for 4 hours. For plasmid transfections, 1 μg of the indicated plasmids and 10 μl of Lipofectamine 2000 reagent were diluted in 100 μl of OptiMEM medium (ThermoFisher). Then, the diluted plasmids were added to diluted Lipofectamine 2000 reagent and incubated for 5 minutes at room temperature. The mixed plasmid-lipid complex was added to each well of a 6-well plate containing cells for 4 hours. The cells were replaced in fresh competed medium after 4 hours transfection.

### Quantification of endosomal imaging data

The automatic computational pipeline for the analysis of endosomal distribution in two-channel TIR-FM imaging data was adapted from our previous work in (Reis et al., 2015) and optimized for high-throughput analysis. The detection algorithm applies the method described in (Aguet et al., 2013). Briefly, the algorithm uses a locally adaptive thresholding approach to obtain a mask of candidate endosome signal followed by subpixellic fitting of a Gaussian function with a scale fixed to 1.5 pixels. Every location presenting intensity significantly above the surrounding local background signal (p-value <0.05) was then classified as an endosome. The cell boundary was estimated based on the EGFP channel after Gaussian smoothing (scale set to 5 pixels). The cell mask is defined as the largest connected component in the set of pixels above a segmentation threshold. The threshold value was estimated for each image using the least probable intensity value lying between the two modes of the histogram of the smoothed image: the first mode describes the background pixel and the second mode describes the cytosolic locations (bin number set to 100). For each cell, the average count-and intensity-measured function of the membrane distance were computed and exported to the Prism (GraphPad) software for flexible representations.

The pipeline has been optimized and parallelized to allow for the processing of hundreds of cell images on their day of acquisition. A summary view of each movie was produced for a systematic review of quantification quality and cellular heterogeneity. The analysis software is available online:

https://cloud.biohpc.swmed.edu/index.php/s/y18NPUBF0YsAXXw

### Thick TIR-FM

Total internal reflection fluorescence microscopy (TIR-FM) was performed as previously described (Loerke et al., 2009). For endosome imaging, we adjusted the illumination of TIRF to provide a theoretical 200nm penetration depth of the evanescent field (therefore named “thick TIRF”) to reduce noise from out of focus planes. Briefly, fixed cells were mounted in PBS and imaged using a 60x, 1.49 NA APO TIRF objective (Nikon) mounted on fully motorized Nikon Ti-Eclipse inverted microscope with Perfect Focus System and coupled to an Andor “Diskovery TIRF/ Borealis widefield illuminator” equipped with an additional 1.8x tube lens (yielding a final magnification of 108x). TIR-FM illumination was achieved using a Diskovery Platform (Andor Technology). During imaging, cells were maintained at 37°C. Image sequences were acquired using a sCMOS camera with 6.5µm pixel size (pco.edge).

### RT-PCR

One 80-90% confluent 6 cm dish of cells was used for RNA extraction by adding 1 ml of TRIzol reagent (Invitrogen). cDNA was prepared from 5 μg of extracted RNA using SuperScript^®^ IV reverse transcriptase (Invitrogen) and oligo dT primers. 3μl of RT product was used for template for human Dyn3 RT-PCR and 1μl of RT product was used to amplify actin. 50μl PCR reactions contained a final concentration of 1.25 U OneTaq^®^ DNA polymerase (BioLab), 200 μM dNTP, 0.2μM of forward and reverse primers, and 1x OneTaq standard reaction buffer. PCR was run for 30 cycles of denaturation (30 seconds at 94°C), annealing (45 seconds), and extension (60 seconds at 68°C).

### Purification and biotinylation of β1 integrin scFv

A single chain variable fragment (scFv) targeting β1 integrin was generated based on the mouse monoclonal K20 and expressed in insect cells (Lakoduk et al., *in preparation).* Briefly, HiFive insect cells obtained from Vincent Tagliabracci (University of Texas Southwestern Medical Center) were infected for 48 hours with P2 baculovirus encoding for the expression of a cell-secreted His_6_-tagged β1 integrin scFv. Insect cell culture supernatant was harvested and purified by FPLC and 1ml HisTrapExcel column (GE Healthcare). Pooled His-purified scFv were further purified via size exclusion chromatography on a Superdex200 Increase (GE Healthcare). Purified recombinant scFv was biotinylated at a 5:1 molar ratio with EZ-Link Sulfo-NHS-SS-Biotin (ThermoFisher) for 2 hours at 4°C for use in biochemical endocytosis and recycling assays. Free biotin was removed using Zeba spin desalting columns (ThermoFisher). Biotinylated scFv (in PBS + 5% glycerol) was flash frozen in liquid nitrogen (LN_2_) in single-use aliquots and stored at -80°C.

### Cell Fractionation

For cell fractionation experiments, 1×10^7^ cells were seeded overnight in duplicate 10cm dishes. After 16 hours, cells were washed 3x with PBS and starved for 1 hour in serum-free biotin-free RPMI. EGF-stimulated samples were treated with 20ng/ml EGF for 10 minutes at 37°C/5%CO_2_. Starved and EGF-stimulated cells were then washed on ice with cold PBS containing 1mM sodium ortho-vandate and lysed directly in ice-cold Cell Fractionation (CF) buffer (250mM sucrose, 20mM HEPES pH 7.5, 1mM EGTA, 2mM MgCl_2_, cOmplete™ protease inhibitor tablet, PhosSTOP™ phosphatase inhibitor tablet, 1mM AEBSF, and 1mM sodium ortho-vandate). Cells were ruptured with three freeze/thaw cycles of freezing in LN_2_ and thawing on ice for 10 minutes. Nuclei and cell debris were cleared by centrifugation at 5,000 rpm for 10 minutes at 4°C. Post-nuclear supernatant was then used as input and membrane and soluble cytoplasm fractions were separated by ultracentrifugation (100,000 x *g* for 1 hour at 4°C). Pelleted fractions were washed in 1x CF buffer and re-pelleted at 100,000 x *g* for 45 minutes at 4°C. Pellets were resuspended in CF buffer to be 10x more concentrated than the input samples. 5X Laemmli buffer was added to input, soluble supernatant, and membrane/pellet fractions to be used for western blotting.

### Dynamin-1 Pull-down

Endogenous dynamin-1 protein levels in H1299 and H1299 cells expressing mutant p53 (R273H) were measured after pull-down using GST-AMPHII_SH3_ (David et al., 1994). Approximately 1×10^7^ cells on 10cm dishes were lysed on ice in pull-down (PD) buffer (20mM HEPES pH 7.4, 150mM KCl, 2mM MgCl_2_, 0.2% Triton X-100, cOmplete™ tablets, PhosSTOP™) via cell scraping. Cell lysates were collected in eppendorf tubes and incubated on ice for 30 minutes, followed by centrifugation at 4°C for 10 minutes at 14,000rpm. Clarified lysate was used to measure protein concentration (A_280_). Lysates were pre-cleared with pre-washed glutathione-Sepharose beads (GE Healthcare) for 30 minutes at 4°C. Approximately 1mg of pre-cleared lysates was incubated with GST-AMPHII_SH3_ bead slurry for one hour at 4°C. Beads were washed 3x in PD buffer at 4°C. Bound endogenous Dynamin 1 was eluted directly from beads in 2x Laemmli buffer (Bio-Rad) and used for western blotting.

### Arf6 Activation Assay

Endogenous Arf6 activation was assessed through a pull-down assay (Cohen and Donaldson, 2010) using the construct pGST-GGA3_VHS-GAT_, which encodes the VHS and ARF-binding domains of GGA3 fused to GST, obtained from Juan Bonifacino (National Institutes of Health). Briefly, 8×10^6^ cells were seeded in duplicate overnight (16h) in biotin-free RPMI in 10cm dishes. The next day, cells were washed 3x with PBS and starved for 1 hour in serum-free biotin-free RPMI. Starved samples were then washed 1x with cold PBS and lysed directly on ice in ice-cold assay buffer (50mM Tris pH 7.5, 100mM NaCl, 2mM MgCl2, 1% Triton X-100, 10% glycerol, cOmplete™ tablets, PhosSTOP™). EGF treated samples were stimulated with 20ng/ml EGF for 10 minutes at 37°C/5%CO_2_. Cells were then washed 1x with cold PBS and lysed on ice in assay buffer. Lysates were immediately subjected to centrifugation at 13,000x *g* for 5 minutes at 4°C. Clarified lysates were incubated with GST-GGA3_VHS-GAT_ beads for 30 minutes at 4°C. Beads were washed on ice 3x in assay buffer. Bound proteins were eluted directly in 2x Laemmli buffer and used for western blotting.

### Western blots of active and total Akt and ERK1/2

10^6^ cells were seeded in triplicate overnight (16h) in biotin-free RPMI on 6cm dishes. The next day, cells were washed 3x with PBS and starved for 1 hour in serum-free biotin-free RPMI. Starved cells were then treated with 20ng/ml of EGF for the indicated time. After PBS wash, cells were immediately lysed in 2x Laemmli buffer (BioRad) and used for western blotting.

### Endocytosis assay

EGFR and β1 integrin internalization experiments were performed using biotinylated-EGF, biotinylated-β1 integrin scFv or K20 β1 integrin antibody respectively. Cells were seeded 6 hours in 96-well plates at a density of 3.5×10^4^ cells/well and incubated with 20 ng/ml of biotinylated-EGF (Invitrogen), or 5μg/ml of biotinylated-β1 integrin scFV or K20 β1 integrin antibody in assay buffer (PBS^4+^: PBS supplemented with 1 mM MgCl_2_, 1 mM CaCl_2_, 5 mM glucose and 1% bovine serum albumin) at 37°C for the indicated time points. Cells were then immediately cooled down (4°C) to stop internalization. The remaining surface-bound biotinylated-EGF, biotinylated-β1 integrin scFv or K20 β1 integrin antibody was removed from the cells by an acid wash step (0.2 M acetic acid, 0.2 M NaCl, pH 2.5). Cells were then washed with cold PBS and then fixed in 4% paraformaldehyde (PFA) (Electron Microscopy Sciences) in PBS for 30 min and further permeabilized with 0.2% Triton X-100/PBS for 10 min. Internalized K20 β1 integrin antibody was assessed using a goat anti-mouse HRP-conjugated antibody (Life Technologies), and internalized biotinylated-EGF or biotinylated-β1 integrin scFv was assessed by streptavidin-POD (Roche). The reaction was further developed with OPD (P1536, Sigma-Aldrich), and then stopped by 5M of H_2_SO_4_. The absorbance was read at 490 nm (Biotek Synergy H1 Hybrid Reader). Internalized ligand was expressed as the percentage of the total surface-bound ligand at 4°C (i.e., without acid wash step), measured in parallel (Reis et al., 2015).

### Recycling assay

EGFR and β1 integrin recycling experiments were performed using biotinylated-EGF or biotinylated-β1 integrin scFv respectively. Cells were seeded 6 hours in 96-well plates at a density of 3×10^4^ cells/well and pulsed with 20 ng/ml of biotinylated-EGF or 5 μg/ml of biotinylated-β1 integrin scFv in PBS^4+^ buffer at 37°C for 5min (for biotinylated-EGF) or 10 min (for biotinylated-scFv). Cells were then immediately cooled down (4°C) to stop internalization. The remaining surface-bound biotinylated-EGF or biotinylated-scFv was cleaved with 10mM TCEP [Tris(2-carboxyethyl)phosphine] for 30 seconds at RT. TCEP reduces the disulfide bond releasing the biotin moiety. Cells were washed with cold PBS^4+^ buffer and then incubated in PBS^4+^ containing 20ng/ml of EGF and 10mM TCEP at 37°C for the indicated time points. Cells were then washed 0.2 M acetic acid/0.2 M NaCl (pH 2.5) and PBS and then fixed in 4% paraformaldehyde (PFA) (Electron Microscopy Sciences) in PBS for 30 min and further permeabilized with 0.2% Triton X-100/PBS for 10 min. Remaining internal biotinylated ligand was assessed by streptavidin-POD (1:10,000 dilution in QPBS, Roche). The reaction was further developed with OPD (P1536, Sigma-Aldrich), and then stopped by 5M of H_2_SO_4_. The absorbance was read at 490 nm (Biotek Synergy H1 Hybrid Reader). The decrease in intracellular biotinylated-EGF of biotinylated-scFv (recycling) was calculated relative to the total pool of ligand internalized during the brief pulse.

### Focal Adhesion Analysis

Focal adhesions were analyzed by both fixed-cell immunofluorescence and live cell microscopy. For fixed-cell immunofluorescence, 3×10^5^ cells were seeded overnight on 22×22mm glass coverslips coated with gelatin (0.02%) and fibronectin (25μg/ml). After 16 hours, cells were washed 3x with PBS and fixed with simultaneous cytoplasm washout in 2% PFA/0.5% Triton X-100 for 2 minutes at RT. Cells were then fixed in 4% PFA for 30 minutes at RT. To block non-specific antibody binding, cells were incubated in Q-PBS for 30 minutes at RT. Cells were incubated with mouse monoclonal anti-paxillin antibody (clone 177, BD Biosciences) diluted 1:250 in Q-PBS for 1 hour at RT. After three PBS washes, the cells were incubated with Alexa Fluor-conjugated secondary antibodies (1:1000 dilution in Q-PBS) at 37°C for 1 hour. Alexa Fluor-conjugated phalloidin was added during the secondary antibody incubation step and used to detect cell boundaries in adhesion quantification analysis. After three PBS washes, the cells were mounted in PBS and imaged by TIR-FM using a 60x, 1.49 NA APO TIRF objective (Nikon) mounted on fully motorized Nikon Ti-Eclipse inverted microscope with Perfect Focus System and coupled to an Andor “Diskovery TIRF/ Borealis widefield illuminator” equipped with an additional 1.8x tube lens (yielding a final magnification of 108x). TIR-FM illumination was achieved using a Diskovery Platform (Andor Technology).

For live cell adhesion analyses, cells were infected with a lentivirus encoding an mRuby2-Paxillin fusion protein under the control of a crippled CMV promoter (Dean et al., 2017). This construct was generated by substituting the mNeonGreen (mNG) from pLVX-CMV100-mNG-Paxillin plasmid, gifted by Kevin Dean (Mohan et al., 2018), for mRuby2 fluorescent protein. H1975 and all engineered cKO cells stably infected with mRuby2-Paxillin were sorted by FACS for very low expression and were used in all live cell adhesion imaging experiments. 3×10^5^ cells were seeded on 22×22mm glass coverslips coated with gelatin (0.02%) and fibronectin (25μg/ml) for 6 hours. Seven minute movies were acquired by TIR-FM with 100ms exposure, at a frame rate of 1 frame per 2 seconds. During imaging, cells were maintained at 37°C. Image sequences were acquired using a sCMOS camera with 6.5µm pixel size (pco.edge).

Focal adhesion analysis was performed using a previously published Focal Adhesion Analysis Package (Han et al., 2015; Mohan et al., 2018). Fixed-cell paxillin immunofluorescence images were used to quantify total cellular adhesion density. Live cell paxillin movies were used to determine adhesion dynamics and lifetimes. The analysis software is available online at https://git.biohpc.swmed.edu/danuser/applications/pipelines/1944.

### Inverted invasion assay

Invasion assays were performed in 96-well dishes (PerkinElmer, Waltham, MA) as previously described (Bendris et al., 2016). In brief, cells were suspended in collagen (1.5 mg/ml) (Nutacon #5409) supplemented with 25μg/ml of human fibronectin (Sigma) at 5×10^4^ cells/ml. 100μl aliquots of collagen/cells were dispensed into the plates. Plates were centrifuged at 1000 rpm at 4°C and then incubated in a 37°C/5% CO2 tissue-culture incubator for 1 hour for polymerization. After that, 30 μl of serum-free medium containing 80 ng/ml of EGF was added on top of the collagen plug. After 48 hrs, cells were fixed/stained with 4% formaldehyde and 10 μg/ml of Hoechst 33342. For quantification, 15 adjacent images were acquired in each well, yielding **∼**90% field of each well. Nuclei labeled with Hoechst 33342 from 0 μm (bottom of the plate) to 200 μm into the collagen plug, with a 30-μm step, were detected with the object counts feature of Nikon Elements. Invasion ratio was calculated as the sum of cell counts at 50, 100, 150 and 2000 μm over cell counts at 0 μm. Results were obtained from at least three independent experiments including four replicates on each day. Bar charts are plotted as mean of all experiments ± SEM.

### TCGA analysis

mRNA expression level data sets of lung cancer (lung adenocarcinoma and squamous cancer carcinoma) were downloaded from National Cancer Institute Genomic Data Commons Data Portal (https://portal.gdc.cancer.gov/, Data Release 10.1 – February 15, 2018). We grouped tumor patients into wild type p53 and mutant p53 groups based on p53 status. Dyn1 RNA expression value read out as Fragments per Kilobase of transcript per Million mapped reads-upper quartile normalization (FPKM-UQ) were used.

### p53 transcription factor binding site (TF-BS) analysis

Prediction of p53 transcription factor binding sites (TF-BS) on 5’ region of DNM1 genomic DNA was conducted using SABiosciences’ proprietary database (http://saweb2.sabiosciences.com/chipqpcrsearch.php?app=TFBS).

## QUANTIFICATION AND STATISTICAL ANALYSIS

### Quantification of fluorescence intensity and number of endosomes

The result was presented as average curve with error bars (shown in Figure 1B, 1C, 2A, 2B, 3D, 4A, 4B, suppl. Figure 2, suppl. Figure 3 and suppl. Figure 5B). Errors bars along each curve represent 95% confidence intervals.

### Analysis of endocytosis and recycling data

Endocytosis and recycling data (Figures 3A, 3C, 3E, and 5A-D) were pooled and averaged from more than three independent experiments and presented as mean±SEM. The statistical significance was analyzed by Student’s t-test. *, p<0.1; **, p<0.05; ***, p<0.01.

### Analysis of RT-PCR and western blot data

RT-PCR and western blots data (Figures 1D, 3F, 4C-F, and suppl. Figure S1B) were pooled and averaged from three independent experiments and presented as mean±SEM. The statistical significance was analyzed by Student’s t-test. *, p<0.1; **, p<0.05; ***, p<0.01.

### Analysis of TCGA RNA expression data

TCGA RNA expression data of NSCLC patients was downloaded from National Cancer Institute Genomic Data Commons Data Portal (https://portal.gdc.cancer.gov/, Data Release 10.1 – February 15, 2018). Pre-stratifying NSCLC patients according to p53 status-wild type p53 and mutant p53 (i.e. deletions and mutations). RNA expression of DNM1 was presented as Box and whisker plots represent the 5-95^th^ percentile (Figure 4H). Mann–Whitney U-test was performed to compute significance; ***P< 0.001.

### Analysis of focal adhesion data

Focal adhesion density and lifetime data was presented as Box and whisker plots represent the 5-95^th^ percentile (Figures 6B-C). Mann–Whitney U-test was performed to compute significance: **P<0.05; ***P<0.001.

### Analysis of invasion data

Invasion data were pooled and averaged from three independent experiments and presented as presented mean ± SEM, n=3. Statistical significance was analyzed by Student’s t test; **P< 0.05, ***P<0.001.

---

**Suppl. Figure 1.**
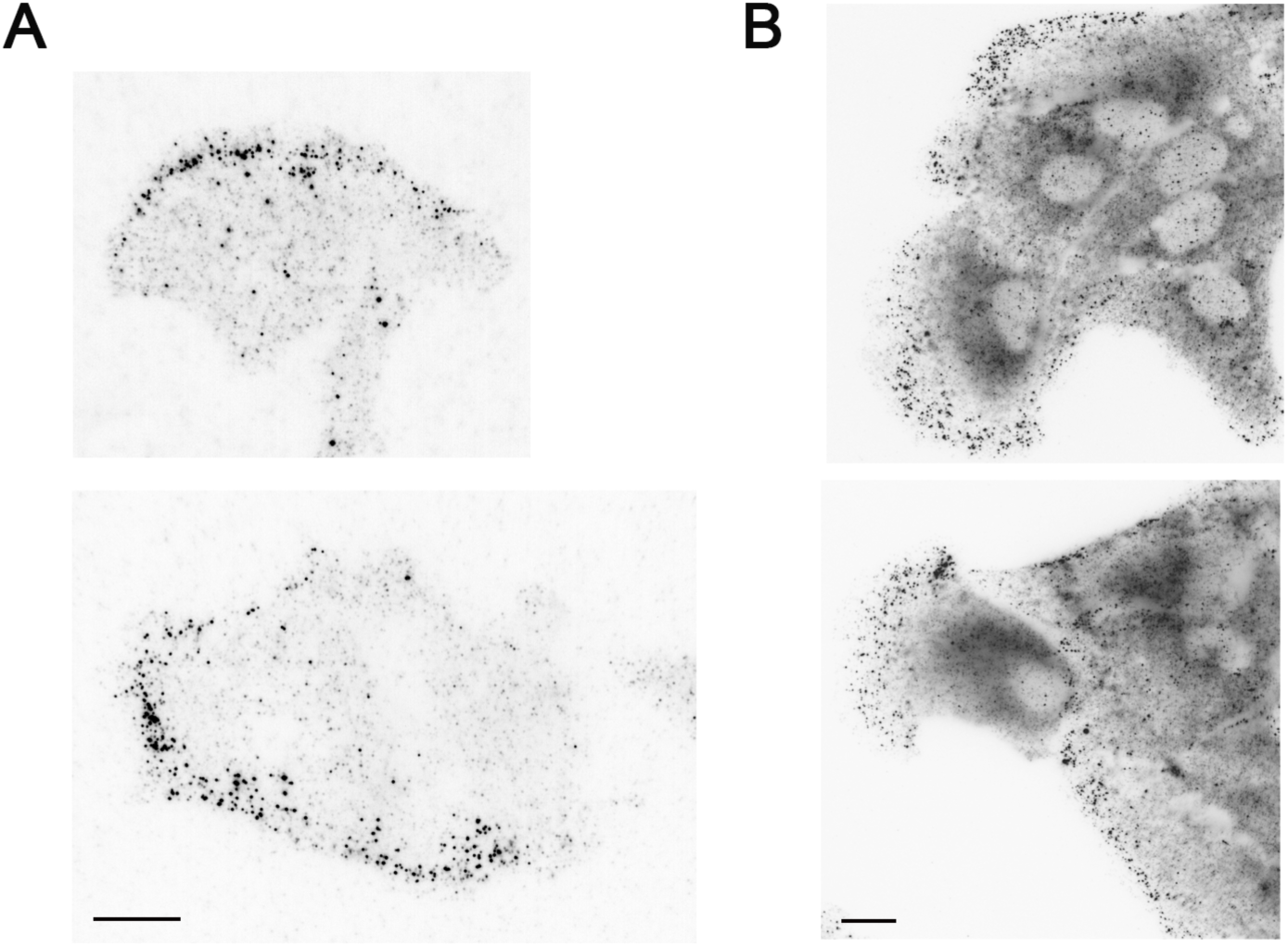
Polarized distribution of perimeter APPL1-positive endosomes in cells. (A) Representative ‘thick-TIRF’ IF images of APPL1 endosome in dispersed H1975 cells. (B) Confocal images of APPL1 endosome in clustered RPE cells. Scale bar, 10μm.

**Suppl. Figure 2.**
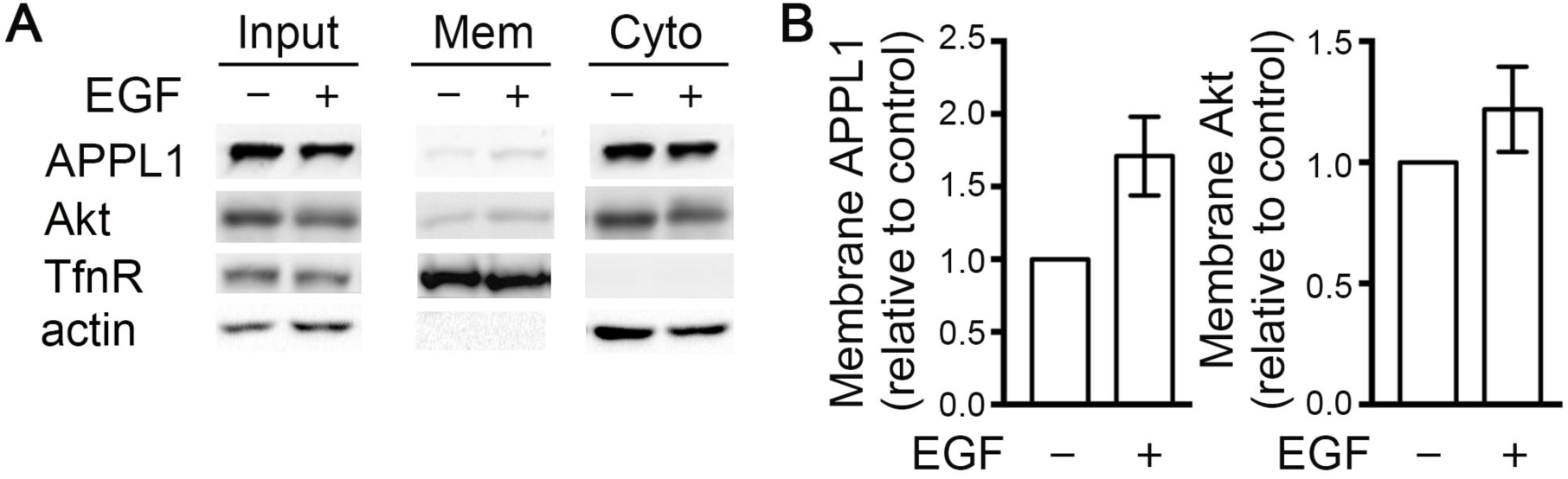
EGF stimulation increases APPL1 membrane recruitment. (A) Representative western blots of cell fractionation of H1975 cells into membrane and cytosolic fractions. TfnR and actin serve as controls for membrane and cytosolic fractions, respectively. Cells were serum-starved and treated, or not, with EGF (20 ng/ml) for 10 minutes. Mem: membrane fraction. Cyto: cytosolic fraction. (B) Quantification of membrane-associated APPL1 and Akt with/without 10 min EGF treatment. n=3.

**Suppl. Figure 3.**
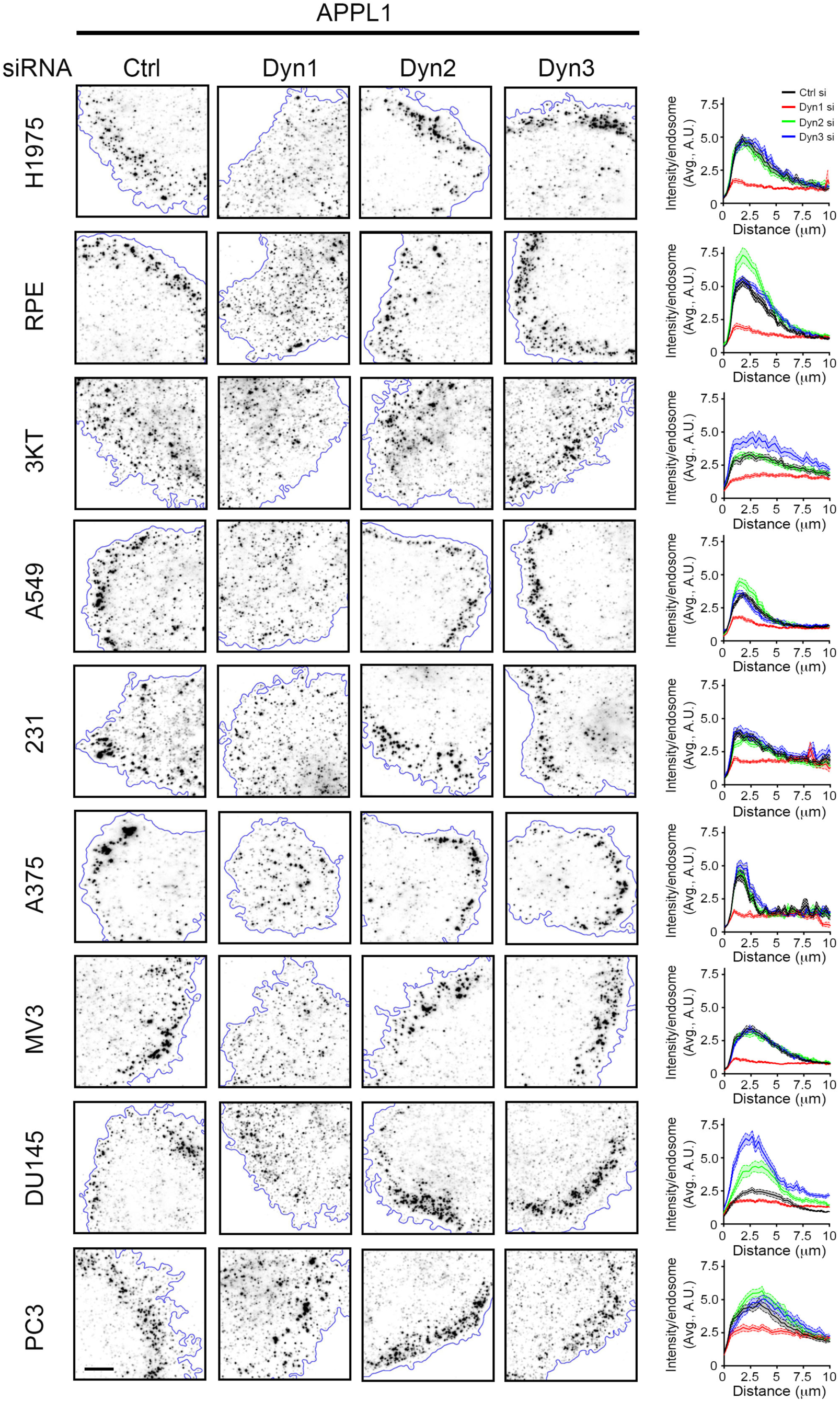
Dynamin-1 is required for EGF-stimulated APPL1 endosome accumulation at the cell edge in all cell lines tested. Representative ‘thick-TIRF’ IF images and accompanying quantification of APPL1 endosome intensity relative to their distance from the cell edge in control (Ctrl), Dyn1, Dyn2 or Dyn3 in the following siRNA treated cell lines (p53 status indicated): H1975 (homozygous p53 R273H), ARPE-19, human retinal pigment epithelia (RPE, WT p53), HBEC-3KT, human bronchial epithelial cells (3KT, WT p53), A549 NSCLC cells (WT p53), MDA-MB-231 breast cancer cells (231, homozygous p53 R280K), melanoma cells A375 and MV3 (both WT p53), prostate cancer cells DU145 (heterozygous P223L and V274F p53) and PC3 (138del p53). Cells were serum-starved and treated with EGF (20 ng/ml) for 10 min prior to fixation. Scale bar, 10μm. Errors bars along each curve represent 95% confidence intervals. n=30 cells/condition.

**Suppl. Figure 4.**
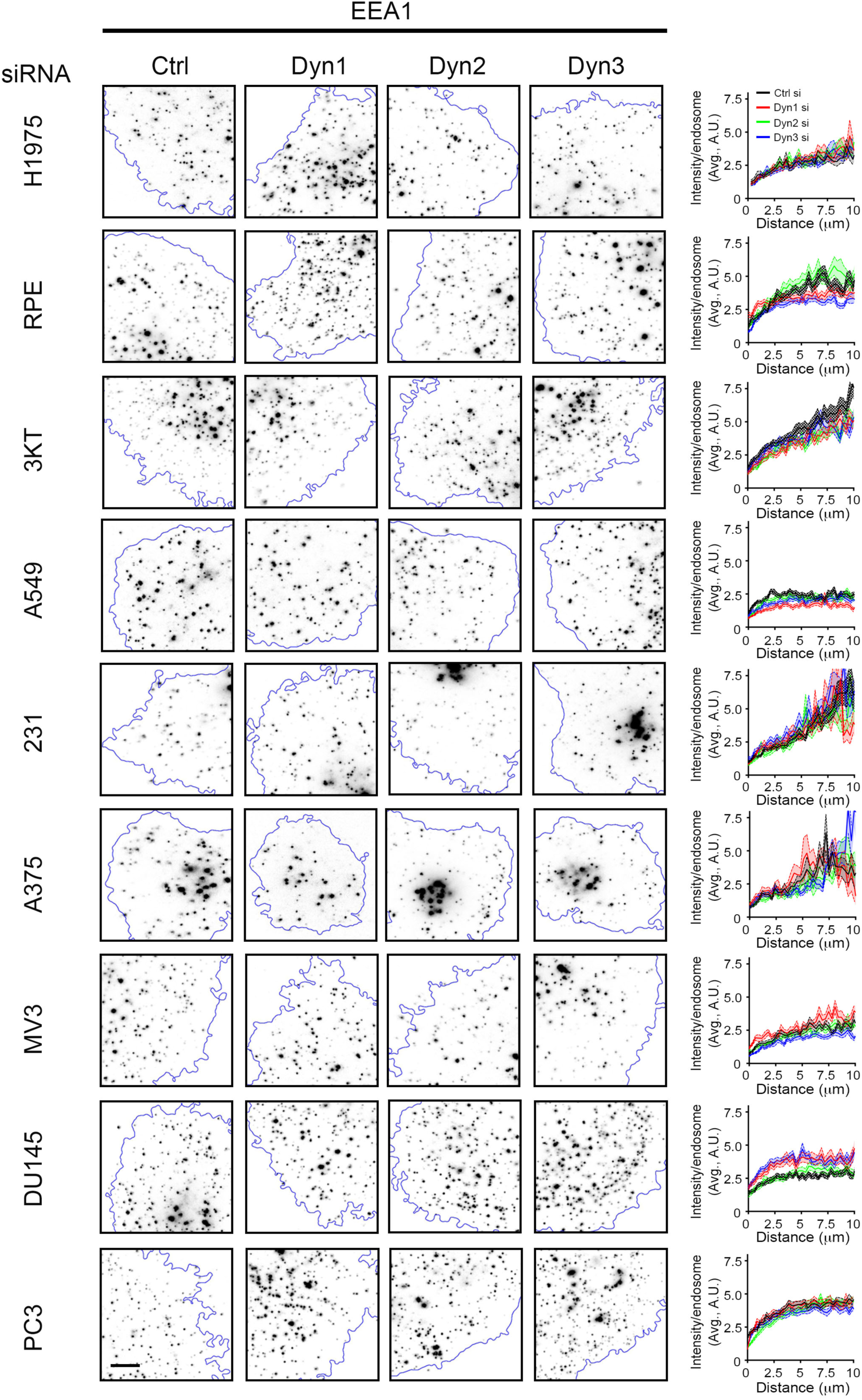
Dynamin-1 does not affect EEA1 endosome distribution. Representative ‘thick-TIRF’ IF images and quantification of EEA1 endosome intensity relative to their distance from the cell edge in control (Ctrl), Dyn1, Dyn2 or Dyn3 siRNA treated cell lines H1975, ARPE-19 (RPE), HBEC-3KT (3KT), A549, MDA-MB-231 (231), A375, MV3, DU145 and PC3. Cell lines are as described in Figure S3. Cells were serum-starved and treated with EGF (20 ng/ml) for 10 min prior to fixation. Scale bar, 10μm. Errors bars along each curve represent 95% confidence intervals. n=30 cells/condition.

**Suppl. Figure 5.**
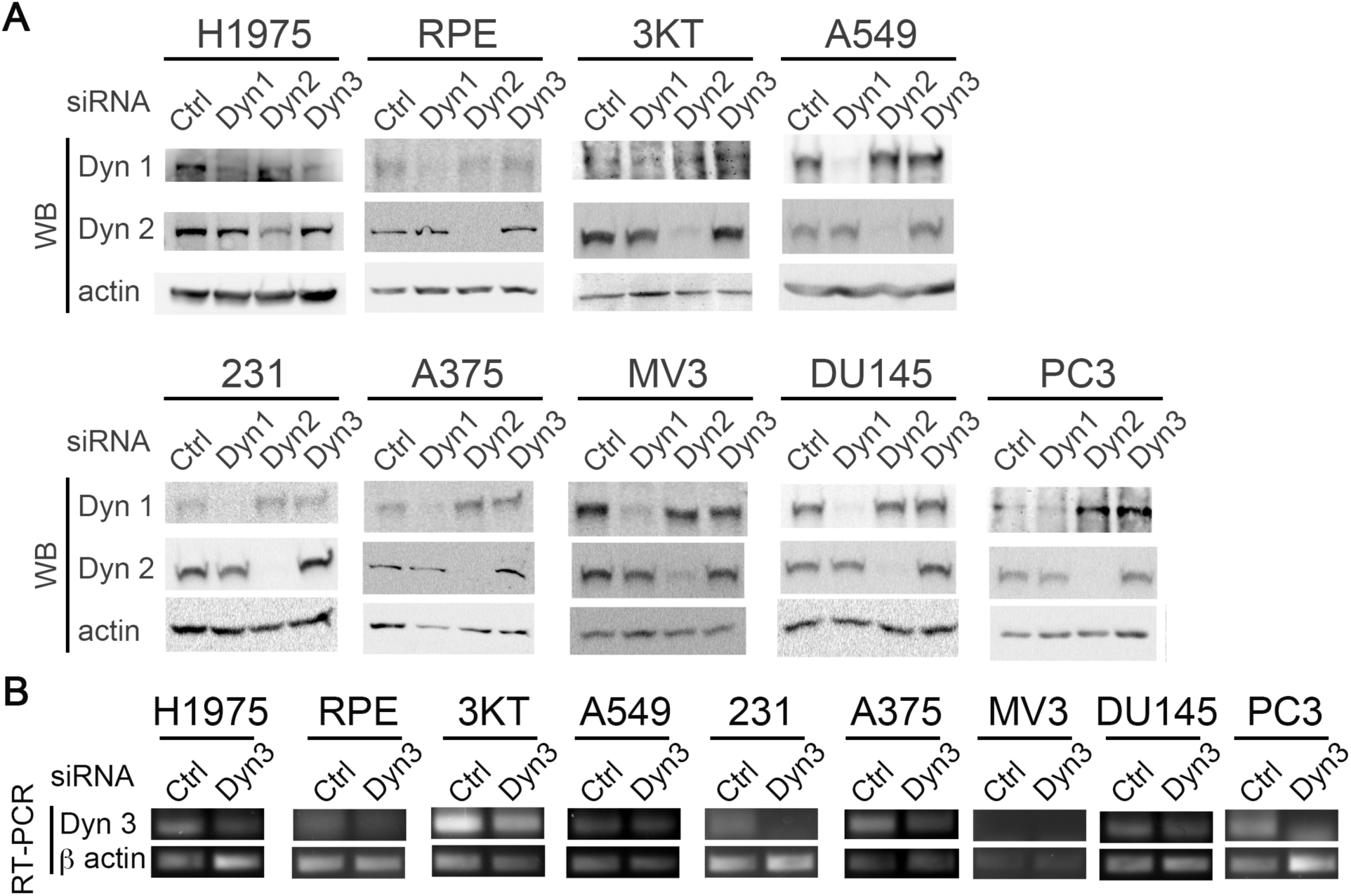
Representative efficiencies of dynamin isoform knockdowns accompanying Figures S2 and S3. Representative western blots of Dyn1 and Dyn2 knockdown efficiency for Suppl. Figures 3-4. (B) Representative RT-PCR of Dyn3 knockdown efficiency for Suppl. Figures 3-4. Actin was used as a loading control in all experiments.

**Suppl. Figure 6.**
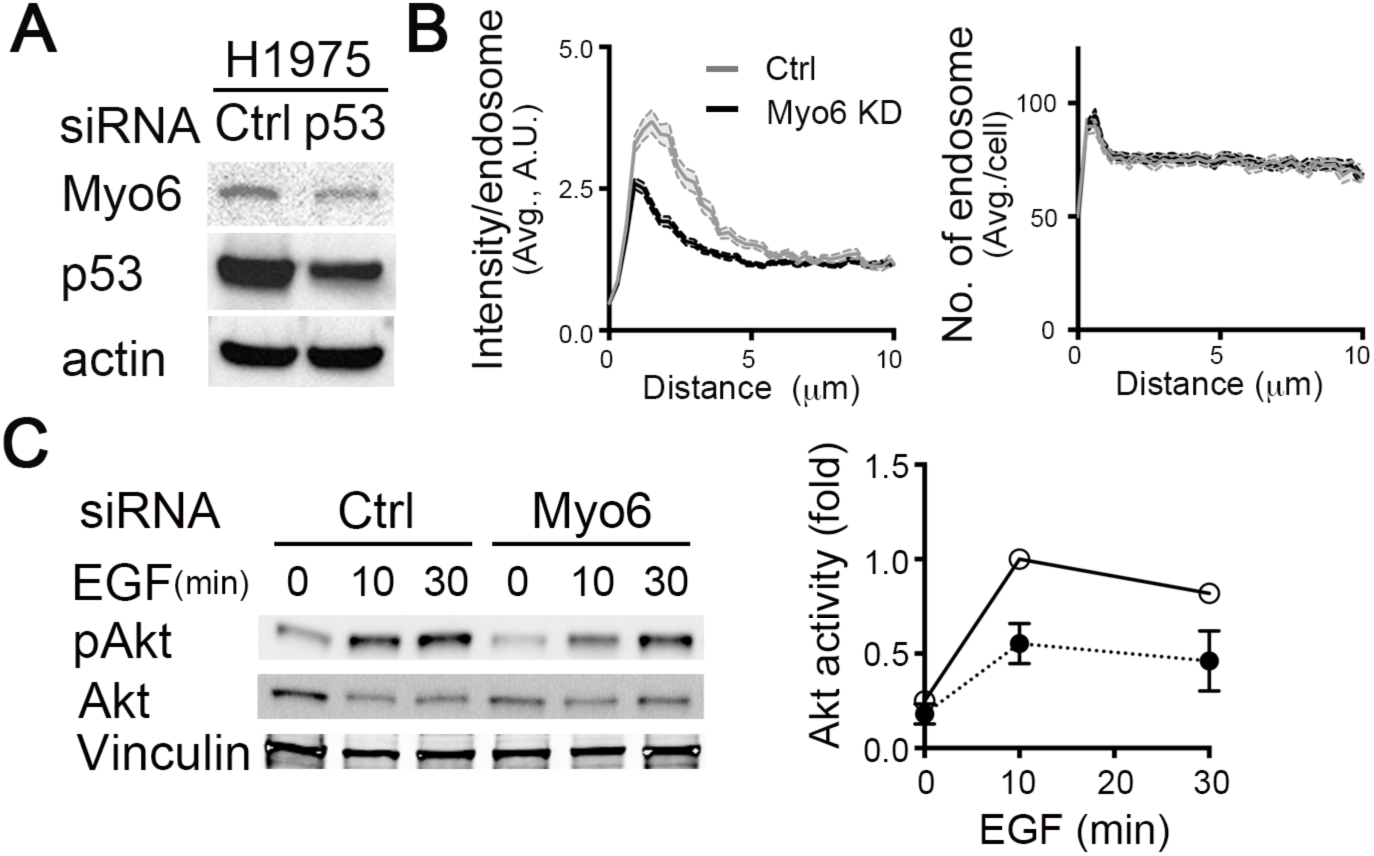
Myo6 is involved mutant p53-driven peripheral APPL1 endosome accumulation and activity. Representative western blots of Myo6 levels in control and p53 KD H1975 cells. Quantification of fluorescence intensity per endosome (left) or number (right) of APPL1-positive endosomes relative to their distance from the cell edge in control or Myo6 KD H1975 cells. Errors bars along each curve represent 95% confidence intervals. n=60 cells/condition. (C) Representative western blots and accompanying quantification of Akt activity in control and Myo6 KD H1975 cells. Cells were serum-starved and treated with EGF (20 ng/ml) for the indicated times before cell lysis. Akt activity was calculated as the ratio of phospho-Akt/total Akt for each time point, then normalized to the control (Ctrl) 10-minute EGF-treated sample.

## References

Aguet, F., Antonescu, C.N., Mettlen, M., Schmid, S.L., and Danuser, G. (2013). Advances in analysis of low signal-to-noise images link dynamin and AP2 to the functions of an endocytic checkpoint. Dev Cell 26, 279–291.

Allaire, P.D., Seyed Sadr, M., Chaineau, M., Seyed Sadr, E., Konefal, S., Fotouhi, M., Maret, D., Ritter, B., Del Maestro, R.F., and McPherson, P.S. (2013). Interplay between Rab35 and Arf6 controls cargo recycling to coordinate cell adhesion and migration. J Cell Sci 126, 722–731.

Amiot, M., Bernard, A., Tran, H.C., Leca, G., Kanellopoulos, J.M., and Boumsell, L. (1986). The human cell surface glycoprotein complex (gp 120,200) recognized by monoclonal antibody K20 is a component binding to phytohaemagglutinin on T cells. Scand J Immunol 23, 109–118.

Arjonen, A., Alanko, J., Veltel, S., and Ivaska, J. (2012). Distinct recycling of active and inactive beta1 integrins. Traffic 13, 610–625.

Bacac, M., and Stamenkovic, I. (2008). Metastatic cancer cell. Annu Rev Pathol 3, 221– 247.

Broussard, J.A., Lin, W.H., Majumdar, D., Anderson, B., Eason, B., Brown, C.M., and Webb, D.J. (2012). The endosomal adaptor protein APPL1 impairs the turnover of leading edge adhesions to regulate cell migration. Mol Biol Cell 23, 1486–1499.

Byron, A., Humphries, J.D., Askari, J.A., Craig, S.E., Mould, A.P., and Humphries, M.J. (2009). Anti-integrin monoclonal antibodies. J Cell Sci 122, 4009–4011.

Caswell, P.T., Chan, M., Lindsay, A.J., McCaffrey, M.W., Boettiger, D., and Norman, J.C. (2008). Rab-coupling protein coordinates recycling of alpha5beta1 integrin and EGFR1 to promote cell migration in 3D microenvironments. J Cell Biol 183, 143–155.

Caswell, P.T., Vadrevu, S., and Norman, J.C. (2009). Integrins: masters and slaves of endocytic transport. Nat Rev Mol Cell Biol 10, 843–853.

Chen, P.H., Bendris, N., Hsiao, Y.J., Reis, C.R., Mettlen, M., Chen, H.Y., Yu, S.L., and Schmid, S.L. (2017). Crosstalk between CLCb/Dyn1-Mediated Adaptive Clathrin-Mediated Endocytosis and Epidermal Growth Factor Receptor Signaling Increases Metastasis. Dev Cell 40, 278–288 e275.

Cohen, L.A., and Donaldson, J.G. (2010). Analysis of Arf GTP-binding protein function in cells. Curr Protoc Cell Biol Chapter 3, Unit 14 12 11–17.

Diggins, N.L., Kang, H., Weaver, A., and Webb, D.J. (2018). alpha5beta1 integrin trafficking and Rac activation are regulated by APPL1 in a Rab5-dependent manner to inhibit cell migration. J Cell Sci 131.

Diggins, N.L., and Webb, D.J. (2017). APPL1 is a multifunctional endosomal signaling adaptor protein. Biochem Soc Trans 45, 771–779.

Ding, Y., Cao, Y., Wang, B., Wang, L., Zhang, Y., Zhang, D., Chen, X., Li, M., and Wang, C. (2016). APPL1-Mediating Leptin Signaling Contributes to Proliferation and Migration of Cancer Cells. PLoS One 11, e0166172.

Fang, Z., Takizawa, N., Wilson, K.A., Smith, T.C., Delprato, A., Davidson, M.W., Lambright, D.G., and Luna, E.J. (2010). The membrane-associated protein, supervillin, accelerates F-actin-dependent rapid integrin recycling and cell motility. Traffic 11, 782–799.

Friedl, P., and Alexander, S. (2011). Cancer invasion and the microenvironment: plasticity and reciprocity. Cell 147, 992–1009.

Goh, L.K., Huang, F., Kim, W., Gygi, S., and Sorkin, A. (2010). Multiple mechanisms collectively regulate clathrin-mediated endocytosis of the epidermal growth factor receptor. J Cell Biol 189, 871–883.

Hsu, P.D., Lander, E.S., and Zhang, F. (2014). Development and applications of CRISPR-Cas9 for genome engineering. Cell 157, 1262–1278.

Ivaska, J., and Heino, J. (2011). Cooperation between integrins and growth factor receptors in signaling and endocytosis. Annu Rev Cell Dev Biol 27, 291–320.

Jean-Alphonse, F., Bowersox, S., Chen, S., Beard, G., Puthenveedu, M.A., and Hanyaloglu, A.C. (2014). Spatially restricted G protein-coupled receptor activity via divergent endocytic compartments. J Biol Chem 289, 3960–3977.

Jones, M.C., Caswell, P.T., and Norman, J.C. (2006). Endocytic recycling pathways: emerging regulators of cell migration. Curr Opin Cell Biol 18, 549–557.

Jung, E.J., Liu, G., Zhou, W., and Chen, X. (2006). Myosin VI is a mediator of the p53-dependent cell survival pathway. Mol Cell Biol 26, 2175–2186.

Kalaidzidis, I., Miaczynska, M., Brewinska-Olchowik, M., Hupalowska, A., Ferguson, C., Parton, R.G., Kalaidzidis, Y., and Zerial, M. (2015). APPL endosomes are not obligatory endocytic intermediates but act as stable cargo-sorting compartments. J Cell Biol 211, 123–144.

Lang, G.A., Iwakuma, T., Suh, Y.A., Liu, G., Rao, V.A., Parant, J.M., Valentin-Vega, Y.A., Terzian, T., Caldwell, L.C., Strong, L.C., et al. (2004). Gain of function of a p53 hot spot mutation in a mouse model of Li-Fraumeni syndrome. Cell 119, 861–872.

Lanzetti, L., and Di Fiore, P.P. (2017). Behind the Scenes: Endo/Exocytosis in the Acquisition of Metastatic Traits. Cancer Res 77, 1813–1817.

Masters, T.A., Tumbarello, D.A., Chibalina, M.V., and Buss, F. (2017). MYO6 Regulates Spatial Organization of Signaling Endosomes Driving AKT Activation and Actin Dynamics. Cell Rep 19, 2088–2101.

Mellman, I., and Yarden, Y. (2013). Endocytosis and cancer. Cold Spring Harb Perspect Biol 5, a016949.

Mitsuuchi, Y., Johnson, S.W., Sonoda, G., Tanno, S., Golemis, E.A., and Testa, J.R. (1999). Identification of a chromosome 3p14.3-21.1 gene, APPL, encoding an adaptor molecule that interacts with the oncoprotein-serine/threonine kinase AKT2. Oncogene 18, 4891–4898.

Muller, P.A., Caswell, P.T., Doyle, B., Iwanicki, M.P., Tan, E.H., Karim, S., Lukashchuk, N., Gillespie, D.A., Ludwig, R.L., Gosselin, P., et al. (2009). Mutant p53 drives invasion by promoting integrin recycling. Cell 139, 1327–1341.

Muller, P.A., Trinidad, A.G., Caswell, P.T., Norman, J.C., and Vousden, K.H. (2014). Mutant p53 regulates Dicer through p63-dependent and-independent mechanisms to promote an invasive phenotype. J Biol Chem 289, 122–132.

Muller, P.A., Trinidad, A.G., Timpson, P., Morton, J.P., Zanivan, S., van den Berghe, P.V., Nixon, C., Karim, S.A., Caswell, P.T., Noll, J.E., et al. (2013). Mutant p53 enhances MET trafficking and signalling to drive cell scattering and invasion. Oncogene 32, 1252–1265.

Olive, K.P., Tuveson, D.A., Ruhe, Z.C., Yin, B., Willis, N.A., Bronson, R.T., Crowley, D., and Jacks, T. (2004). Mutant p53 gain of function in two mouse models of Li-Fraumeni syndrome. Cell 119, 847–860.

Onodera, Y., Nam, J.M., Hashimoto, A., Norman, J.C., Shirato, H., Hashimoto, S., and Sabe, H. (2012). Rab5c promotes AMAP1-PRKD2 complex formation to enhance beta1 integrin recycling in EGF-induced cancer invasion. J Cell Biol 197, 983–996.

Paul, N.R., Jacquemet, G., and Caswell, P.T. (2015). Endocytic Trafficking of Integrins in Cell Migration. Curr Biol 25, R1092–1105.

Reis, C.R., Chen, P.H., Bendris, N., and Schmid, S.L. (2017). TRAIL-death receptor endocytosis and apoptosis are selectively regulated by dynamin-1 activation. Proc Natl Acad Sci U S A 114, 504–509.

Reis, C.R., Chen, P.H., Srinivasan, S., Aguet, F., Mettlen, M., and Schmid, S.L. (2015). Crosstalk between Akt/GSK3beta signaling and dynamin-1 regulates clathrin-mediated endocytosis. EMBO J 34, 2132–2146.

Roberts, M., Barry, S., Woods, A., van der Sluijs, P., and Norman, J. (2001). PDGF-regulated rab4-dependent recycling of alphavbeta3 integrin from early endosomes is necessary for cell adhesion and spreading. Curr Biol 11, 1392–1402.

Schenck, A., Goto-Silva, L., Collinet, C., Rhinn, M., Giner, A., Habermann, B., Brand, M., and Zerial, M. (2008). The endosomal protein Appl1 mediates Akt substrate specificity and cell survival in vertebrate development. Cell 133, 486–497.

Schmid, S.L. (2017). Reciprocal regulation of signaling and endocytosis: Implications for the evolving cancer cell. J Cell Biol 216, 2623–2632.

Senturk, S., Shirole, N.H., Nowak, D.G., Corbo, V., Pal, D., Vaughan, A., Tuveson, D.A., Trotman, L.C., Kinney, J.B., and Sordella, R. (2017). Rapid and tunable method to temporally control gene editing based on conditional Cas9 stabilization. Nat Commun 8, 14370.

Setodji, C.M., McCaffrey, D.F., Burgette, L.F., Almirall, D., and Griffin, B.A. (2017). The Right Tool for the Job: Choosing Between Covariate-balancing and Generalized Boosted Model Propensity Scores. Epidemiology 28, 802–811.

Sever, R., and Brugge, J.S. (2015). Signal transduction in cancer. Cold Spring Harb Perspect Med 5.

Sposini, S., Jean-Alphonse, F.G., Ayoub, M.A., Oqua, A., West, C., Lavery, S., Brosens, J.J., Reiter, E., and Hanyaloglu, A.C. (2017). Integration of GPCR Signaling and Sorting from Very Early Endosomes via Opposing APPL1 Mechanisms. Cell Rep 21, 2855–2867.

Srinivasan, S., Burckhardt, C.J., Bhave, M., Chen, Z., Chen, P.H., Wang, X., Danuser, G., and Schmid, S.L. (2018). A noncanonical role for dynamin-1 in regulating early stages of clathrin-mediated endocytosis in non-neuronal cells. PLoS Biol 16, e2005377.

Takada, Y., and Puzon, W. (1993). Identification of a regulatory region of integrin beta 1 subunit using activating and inhibiting antibodies. J Biol Chem 268, 17597–17601.

Tan, Y., Xin, X., Coffey, F.J., Wiest, D.L., Dong, L.Q., and Testa, J.R. (2016). Appl1 and Appl2 are Expendable for Mouse Development But Are Essential for HGF-Induced Akt Activation and Migration in Mouse Embryonic Fibroblasts. J Cell Physiol 231, 1142–1150.

Tan, Y., You, H., Wu, C., Altomare, D.A., and Testa, J.R. (2010). Appl1 is dispensable for mouse development, and loss of Appl1 has growth factor-selective effects on Akt signaling in murine embryonic fibroblasts. J Biol Chem 285, 6377–6389.

Tomas, A., Futter, C.E., and Eden, E.R. (2014). EGF receptor trafficking: consequences for signaling and cancer. Trends Cell Biol 24, 26–34.

Wandinger-Ness, A., and Zerial, M. (2014). Rab proteins and the compartmentalization of the endosomal system. Cold Spring Harb Perspect Biol 6, a022616.

Weissman, A.M., Klausner, R.D., Rao, K., and Harford, J.B. (1986). Exposure of K562 cells to anti-receptor monoclonal antibody OKT9 results in rapid redistribution and enhanced degradation of the transferrin receptor. J Cell Biol 102, 951–958.

Zoncu, R., Perera, R.M., Balkin, D.M., Pirruccello, M., Toomre, D., and De Camilli, P. (2009). A phosphoinositide switch controls the maturation and signaling properties of APPL endosomes. Cell 136, 1110–1121.

## Reference

Bendris, N., Williams, K.C., Reis, C.R., Welf, E.S., Chen, P.H., Lemmers, B., Hahne, M., Leong, H.S., and Schmid, S.L. (2016). SNX9 promotes metastasis by enhancing cancer cell invasion via differential regulation of RhoGTPases. Mol Biol Cell.27, 1409–1551

Cohen, L.A., and Donaldson, J.G. (2010). Analysis of Arf GTP-binding protein function in cells. Curr Protoc Cell Biol Chapter 3, Unit 14 12 11-17.

David, C., Solimena, M., and De Camilli, P. (1994). Autoimmunity in stiff-Man syndrome with breast cancer is targeted to the C-terminal region of human amphiphysin, a protein similar to the yeast proteins, Rvs167 and Rvs161. FEBS Lett 351, 73–79.

Dean, K.M., Roudot, P., Welf, E.S., Pohlkamp, T., Garrelts, G., Herz, J., and Fiolka, R. (2017). Imaging Subcellular Dynamics with Fast and Light-Efficient Volumetrically Parallelized Microscopy. Optica 4, 263–271.

Han, S.J., Oak, Y., Groisman, A., and Danuser, G. (2015). Traction microscopy to identify force modulation in subresolution adhesions. Nat Methods 12, 653–656.

Loerke, D., Mettlen, M., Yarar, D., Jaqaman, K., Jaqaman, H., Danuser, G., and Schmid, S.L. (2009). Cargo and dynamin regulate clathrin-coated pit maturation. PLoS Biol 7, e57.

Mohan, A.S., Dean, K.S., Kasitinon, S.Y., Isogai, T., Murali, V.S., Han, S.J., Roudot, P., Groisman, A., Welf, E., and Danuser, G. (2018). Hyperactive Rac1 drives MAPK-independent proliferation in melanoma by assembly of a mechanosensitive dendritic actin network. BioRxiv. doi: https://doi.org/10.1101/326710

